# Oligomerization and positive feedback on membrane recruitment encode dynamically stable PAR-3 asymmetries in the *C. elegans* zygote

**DOI:** 10.1101/2023.08.04.552031

**Authors:** Charles F Lang, Ondrej Maxian, Alexander Anneken, Edwin Munro

## Abstract

Studies of PAR polarity have emphasized a paradigm in which mutually antagonistic PAR proteins form complementary polar domains in response to transient cues. A growing body of work suggests that the oligomeric scaffold PAR-3 can form unipolar asymmetries without mutual antagonism, but how it does so is largely unknown. Here we combine single molecule analysis and modeling to show how the interplay of two positive feedback loops promotes dynamically stable unipolar PAR-3 asymmetries in early *C. elegans* embryos. First, the intrinsic dynamics of PAR-3 membrane binding and oligomerization encode negative feedback on PAR-3 dissociation. Second, membrane-bound PAR-3 promotes its own recruitment through a mechanism that requires the anterior polarity proteins PAR-6 and PKC-3. Using a kinetic model tightly constrained by our experimental measurements, we show that these two feedback loops are individually required and jointly sufficient to encode dynamically stable and locally inducible unipolar PAR-3 asymmetries in the absence of posterior inhibition. Given the central role of PAR-3, and the conservation of PAR-3 membrane-binding, oligomerization, and core interactions with PAR-6/PKC-3, these results have widespread implications for PAR-mediated polarity in metazoa.

## Introduction

The PAR proteins are a highly conserved network of proteins that govern cell polarity in many different cells and tissues across the metazoa^1,2^. During polarization, the PAR proteins become asymmetrically enriched in response to transient local cues. These asymmetries are reinforced through feedback loops in which PAR proteins locally promote or inhibit one another’s accumulation. Asymmetrically enriched PARs then act through different downstream targets to elaborate functionally polarized states. While the local polarizing cues vary widely across different contexts, the interactions among PAR proteins that govern the response to these cues is more highly conserved. Thus, a central challenge is to understand how systems of feedback among the PARs encode their ability to form and stabilize spatial patterns of enrichment that define functional polarity.

The one-cell *C. elegans* embryo (i.e., the zygote) has been a powerful model system for uncovering the core molecular circuitry that governs PAR polarity. During polarization of the zygote, two sets of PAR proteins become asymmetrically enriched in complementary anterior and posterior membrane domains. The anterior aPARs include the oligomeric scaffold PAR-3, the adaptor PAR-6, the kinase PKC-3, and the small GTPase CDC-42. The posterior pPARs include the kinase PAR-1, a *C. elegans*-specific protein called PAR-2, and a putative CDC-42 GAP called CHIN-1.

Polarization proceeds through two distinct phases, called establishment and maintenance^3^, which correspond respectively to mitotic interphase and mitosis. Before polarization, aPARs are uniformly enriched at the cell surface, while pPARs are largely cytoplasmic. During polarity establishment, a local sperm-derived cue triggers the redistribution of aPARs and pPARs into complementary domains. The sperm cue is closely associated with a centrosomal microtubule organizing center called the sperm MTOC that forms near the site of sperm entry ^4,5^. The sperm MTOC acts through the aurora kinase AIR-1 ^6–9^ to trigger actomyosin-based cortical flows that transport aPARs towards the anterior pole ^10,11^. In addition, astral microtubules associated with the sperm MTOC promote the local accumulation of pPARs PAR-1 and PAR-2 on the posterior membrane ^12,13^.

During mitosis, the centrosomes (thus the polarity cue) migrate to the cell center. However, PAR asymmetries are maintained through mitosis despite continuous diffusion and exchange of PAR proteins between the cytoplasm and the cell membrane ^14,15^. The prevailing model is that PAR asymmetries are maintained by a network of mutually antagonistic interactions in which aPARs and pPARs act locally to inhibit one another’s accumulation (reviewed in Lang and Munro ^2^, Figure 1A). Both PAR-3 and active CDC-42 are required for local recruitment of PAR-6/PKC-3 into an active complex with CDC-42 ^16,17^. Active PKC-3 inhibits local accumulation of pPARs, including PAR-1 and CHIN-1 ^12,17–22^. PAR-1 inhibits local accumulation of PAR-3 ^12,23^, while CHIN-1 inhibits local activation of CDC-42 ^17,22,24^. Because both PAR-3 and CDC-42 are required for local recruitment of PAR-6/PKC-3 during mitosis, either PAR-1 or CHIN-1 is sufficient to prevent posterior recruitment of PAR-6/PKC-3.

**Figure 1.**
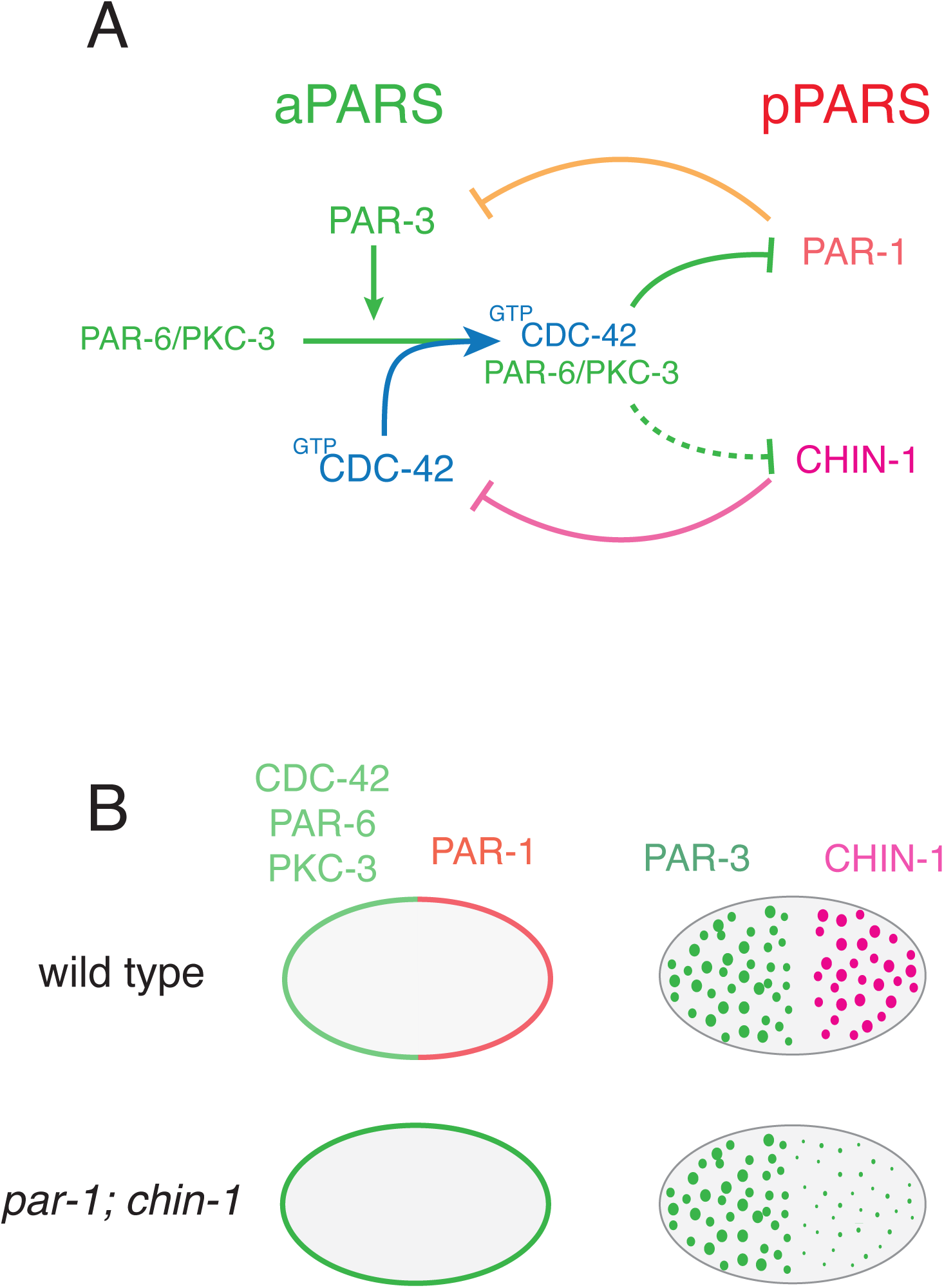
PAR polarity in the *C. elegans* embryo. **(A)** Schematic illustrating key interactions that maintain PAR asymmetries in the *C. elegans* zygote. On the left, PAR-3 promotes entry of the PAR-6/PKC-3 dimer into a complex with active CDC-42. Within this complex, PKC-3 inhibits cortical accumulation of posterior PARs. PAR-1 inhibits cortical accumulation of PAR-3, while CHIN-1 inhibits CDC-42 activity. **(B)** In embryos lacking posterior inhibitors PAR-1 and CHIN-1, anterior PARs CDC-42, PAR-6 and PKC-3 are uniformly distributed at the membrane, and all other posterior PARs are uniformly cytoplasmic. Nevertheless, PAR-3 oligomers remain asymmetrically distributed.

Strikingly, in zygotes that lack both PAR-1 and CHIN-1, all other pPARS are restricted to the cytoplasm, the aPARs CDC-42, PAR-6 and PKC-3 are uniformly enriched on the cell surface, but PAR-3 asymmetries still form and persist through mitosis ^17^ (Figure 1B). Thus, additional mechanisms must exist to maintain unipolar PAR-3 asymmetries in the absence of mutual antagonism with pPARs. PAR-3 is the keystone member of the PAR network without which all other PAR asymmetries are lost, and segregation of PAR-3 is a first step to polarization in many different contexts ^12,25–28^. Therefore, the mechanisms that sustain unipolar PAR-3 asymmetries in embryos that lack posterior inhibition must also contribute to shaping PAR asymmetries in normal embryos. Importantly, unipolar PAR-3 asymmetries have also been observed under wild-type conditions in other contexts, including in neuroblast stem cells ^29–31^, in mammalian hippocampal neurons ^32^ and in *Drosophila* male germline cells ^33^. However, the underlying molecular mechanisms remain unknown.

A key feature of PAR-3 is its ability to form oligomers at the cell membrane. PAR-3 self-associates through a conserved N-terminal PB1-like domain, called the CR1 domain ^34^. The purified CR1 domain assembles head to tail into helical filaments *in vitro* ^35,36^. While PAR-3 contains multiple domains that can support direct binding to plasma membranes ^37,38^, deletion or mutation of the CR1 domain prevents strong accumulation of PAR-3 in *C. elegans* ^37^ and other systems ^34,35,38^, and induces defects in embryonic polarity ^16,37,39^. Oligomerization of PAR-3 plays an essential role in polarity establishment by coupling PAR-3 to the actomyosin cortex, facilitating segregation of PAR-3 and other aPARs by anterior-directed cortical actomyosin flows ^16,26,39–41^. But it remains unknown whether oligomerization of PAR-3 plays additional roles in polarity maintenance, and how oligomerization of PAR-3 might contribute to stabilizing PAR-3 asymmetries in the absence of mutual antagonism among the PARs.

Here, we combine single molecule imaging and particle tracking analysis with genetic manipulations and modeling to reveal a mechanism for self-stabilizing PAR-3 asymmetries in the *C. elegans* zygote. We show that PAR-3 asymmetries are maintained despite the continuous rapid exchange of subunits within PAR-3 oligomers. We find that two key factors maintain PAR-3 asymmetries in the absence of posterior inhibition: First, size-dependent membrane binding avidity of PAR-3 oligomers confers negative dependence of PAR-3 dissociation rate on local PAR-3 density – a form of positive feedback. Second, enhanced recruitment of PAR-3 monomers to membranes where PAR-3 is already enriched, which requires the aPARs CDC-42, PAR-6 and PKC-3, provides a second form of positive feedback. Mathematical models, tightly constrained by our experimental measurements, show that the combination of positive feedback through size-dependent binding avidity and positive feedback on monomer recruitment ensures the coexistence of two dynamically stable states: an unpolarized state, characterized by uniformly high levels of PAR-3, and a polarized state, characterized by asymmetric enrichment of PAR-3. The model predicts that transient local depletion of PAR-3, as occurs in response to the sperm cue during polarity establishment, can induce a transition between unpolarized and polarized states. Thus, self-stabilizing PAR-3 asymmetries encode a lasting memory of PAR distributions induced by a transient polarizing cue. Together, these results reveal a novel mechanism for stabilizing PAR-3 asymmetries in the absence of mutual antagonism, which depends on core dynamics of PAR-3 oligomer assembly and interactions with its conserved binding partners PAR-6 and PKC-3. We propose that similar mechanisms may underlie the stabilization of unipolar PAR asymmetries in other contexts.

## Results

### PAR-3 asymmetries approach a dynamic steady state during maintenance phase

We previously showed that PAR-3 asymmetries persist during maintenance phase in embryos co-depleted of the posterior inhibitors PAR-1 and CHIN-1, when all other known PAR asymmetries are absent ^17^. PAR-1 acts directly by phosphorylating PAR-3 to inhibit its accumulation ^12,37^, while mutating *chin-1* has no further effect on PAR-3 asymmetries produced by depleting PAR-1 ^17^. Thus, we focused here on determining the mechanisms that maintain unipolar PAR-3 asymmetries in the absence of PAR-1. As a first step we used near-TIRF microscopy and single particle analysis to quantify dynamic changes in PAR-3 oligomer size and density during polarization in *par-1* mutant embryos. Consistent with previous reports, mean oligomer size, oligomer density, and overall density of PAR-3 increased steadily on the anterior cortex and decreased on the posterior cortex during polarity establishment, as cortical flows transport PAR-3 towards the anterior pole (Figure 2A-D) ^39,41^. At the onset of maintenance phase, mean oligomer size and overall density of PAR-3 began to decrease, while oligomer density remained roughly constant. The rate of decrease was initially fast but slowed progressively during maintenance phase, such that oligomer size and overall density approached plateaus at ∼50% of their maximum values in late establishment phase. We observed similar PAR-3 oligomer dynamics in control (*par-1* heterozygote) embryos (Figure 2E-H) and throughout the cortex in embryos depleted of SPD-5 by RNAi, which lack a functional sperm cue and fail to establish polarity (Figure S1, Movie S1).

**Figure 2.**
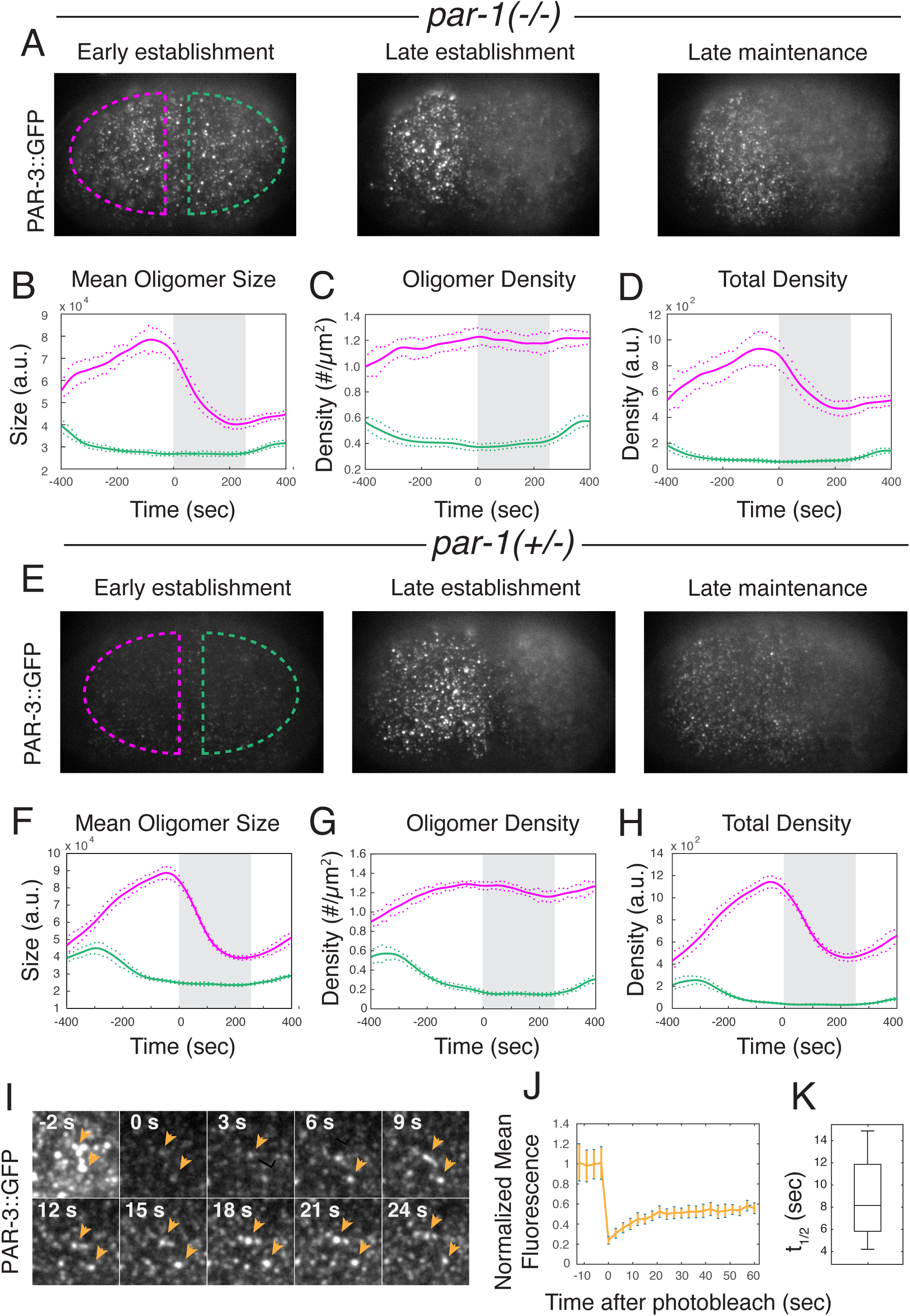
PAR-3 asymmetries approach a steady state during mitosis despite rapid continuous turnover. **(A)** Individual near-TIRF images from a time-lapse sequence showing PAR-3 oligomer intensities and distributions at the cell surface in homozygous *par-1* mutant embryos at early establishment, late establishment and late maintenance phase. **(B-D)** Results of particle detection analysis showing **(B)** mean oligomer size, **(C)** oligomer density, and **(D)** total PAR-3 fluorescence density over time on the anterior (magenta) and posterior (blue) membrane in homozygous *par-1-/-* mutant embryos. Data are from n = 12 embryos aligned with respect to the onset of maintenance phase. The grey shaded area indicates maintenance phase. Solid lines indicate the mean and dots indicate S.E.M. at each time point. **(E)** Individual near-TIRF images from a time-lapse sequence showing PAR-3 oligomer intensities and distributions at the cell surface in control (*par-1+/-* heterozygote) embryos at early establishment, late establishment and late maintenance phase. **(F-H)** Results of particle detection analysis showing **(F)** mean oligomer size, **(G)** oligomer density, and **(H)** total PAR-3 fluorescence density over time on the anterior (magenta) and posterior (blue) cortex in control embryos. Data are from n = 8 embryos aligned with respect to the onset of maintenance phase. The grey shaded area indicates maintenance phase. Solid lines indicate the mean and dots indicate SEM. at each time point. **(I)** Representative micrographs showing the time course of recovery of PAR-3::GFP in the anterior domain after photobleaching the entire field of view in an embryo in early maintenance phase. Timestamps indicate seconds after photobleaching. Orange arrowheads indicate individual PAR-3 oligomers that persist during recovery. **(J)** Plots of average fluorescence density vs time before and after photobleaching, normalized to prebleach levels and corrected for the overall decrease in fluorescence observed during maintenance phase (see Methods). Time zero coincides with the end of the photobleaching pulse. Error bars indicate SEM (n = 4 embryos). **(K)** Box plot showing time to half maximal recovery (t1/2) after photobleaching.

The slow approach to plateau levels of PAR-3 density and mean oligomer size could reflect either (a) the gradual disassembly of otherwise stable PAR-3 oligomers, or (b) the approach to a steady state governed by a dynamic balance of more rapid membrane-binding and oligomer-assembly kinetics. Consistent with the latter possibility, after photobleaching the entire cortex, total fluorescence in detected PAR-3 oligomers recovered to ∼ 50% of the pre-bleach value with a mean half-time to 50% recovery of 8.52 +\- 4.3 seconds (Figure 2I-K; n = 4 embryos; Movie S2). Individual PAR-3 oligomers recovered fluorescence on similarly fast timescales (orange arrowheads in Fig 2I). These data suggest that PAR-3 asymmetries approach a steady state that is maintained in the face of more rapid exchange of cytoplasmic PAR-3 with the membrane and with membrane-bound oligomers.

### A simple model for PAR-3 membrane binding and oligomerization

To probe the mechanisms that underlie this dynamic steady state, we used single molecule imaging and particle tracking analysis to characterize how the kinetics of PAR-3 membrane binding and oligomerization shape oligomer sizes, densities and dissociation rates during late maintenance phase. As a framework for this analysis, we considered a simple kinetic model (Figure 3A, Modeling Supplement) in which: (a) cytoplasmic PAR-3 monomers bind reversibly to the plasma membrane with rate constants *k_on_* and *k_off_*, (b) membrane-bound PAR-3 monomers bind reversibly to one another and to existing oligomers with rate constants *k_ass_* and *k_diss_*, and thus with self-affinity 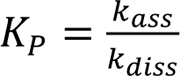, and (c) oligomers dissociate from the membrane at a rate *k_off_*(*n*) that depends on oligomer size n. We initially assumed (and confirm experimentally below) that direct binding of cytoplasmic monomers to PAR-3 oligomers is slow relative to membrane binding, and can therefore be neglected.

**Figure 3.**
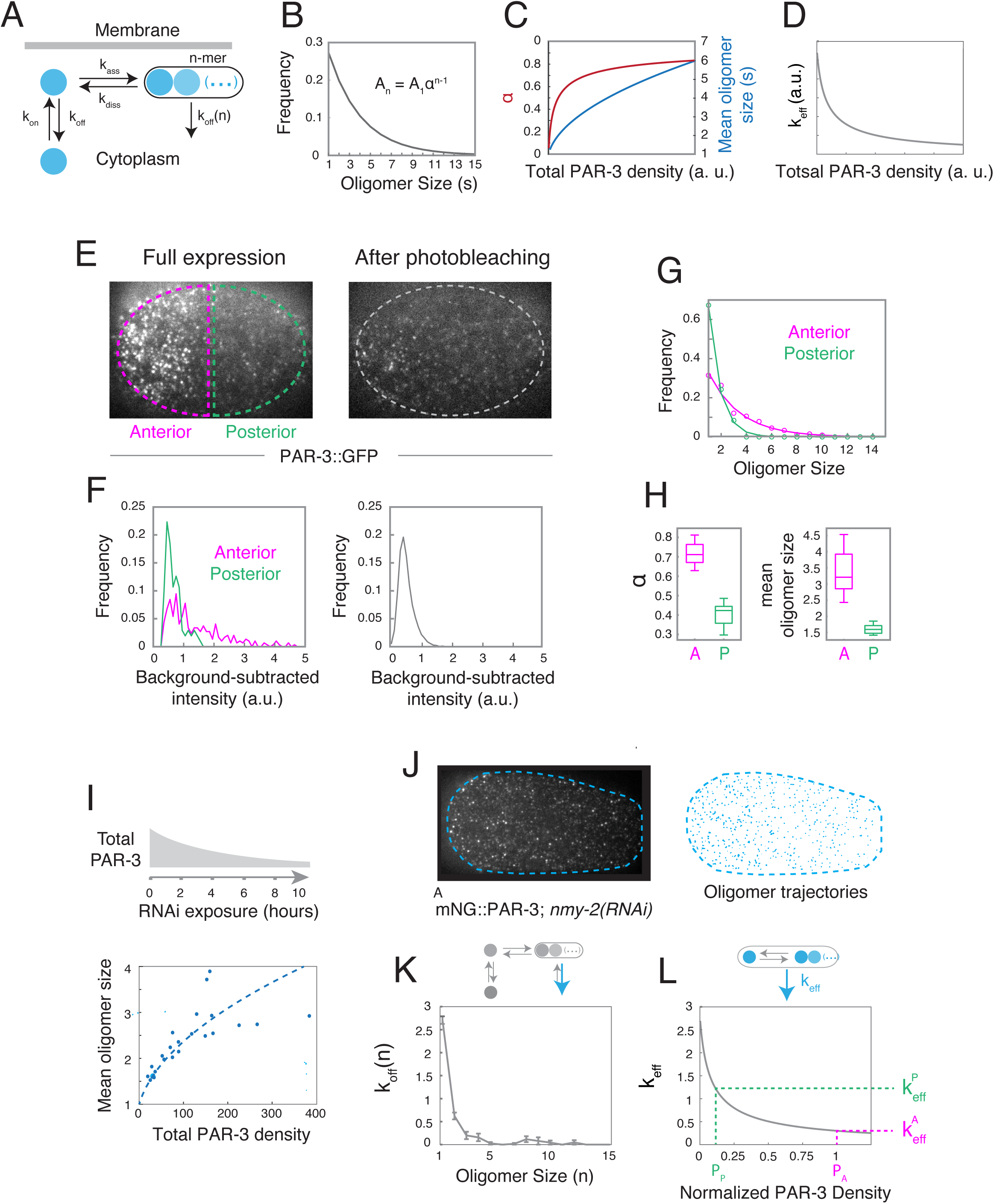
PAR-3 oligomer size distributions reflect kinetics of membrane binding and polymerization. **(A)** A simple kinetic model for PAR-3 oligomerization dynamics. Monomers bind reversibly to the membrane with rate constants *k_on_*and *k_off_*. Membrane-bound monomers bind oligomers with rate constants *k_ass_* and *k_diss_*, and oligomers dissociate from the membrane with size-dependent rate constant *k_off_*(*n*) where *n* is the number of subunits in the oligomer. We neglect direct binding of cytoplasmic monomers to membrane bound oligomers and membrane binding of oligomers with size n > 1 (see text for justification). **(B)** Predicted steady state distribution of oligomer sizes *A*_*n*_ = *A*_1_α^*n*–1^ (see text for details) **(C)** Predicted steady state dependence of oligomerization strength (α) and mean oligomer size (s) on PAR-3 density. **(D)** Predicted steady state dependence of effective dissociation rate *k_eff_* on PAR-3 density. **(E)** Micrographs showing PAR-3 oligomers before (left panel) and after 30 seconds of continuous laser exposure at full power (right panel) in late maintenance phase *par-1* mutant embryos. Dashed orange and green outline the regions in which anterior and posterior measurements were made. **(F) (left)** Distributions of anterior (orange) and posterior (green) speckle intensities measured in a representative embryo. **(right)** Distributions of speckle intensities after photobleaching (gray) in the same embryo. **(G)** Inferred distributions of PAR-3 oligomer sizes on the anterior and posterior cortex of a representative embryo, based on calibrating raw intensities to the estimated intensity distribution of single molecules. Dashed lines indicate data fit to the equation *A*_*n*_ = *A*_1_α^*n*–1^. **(H)** Estimates of oligomerization strength α (left) and mean oligomer size s (right) on the anterior and posterior of n = 8 individual embryos in late maintenance phase. **(I) (top)** Total pool of PAR-3 can be tuned by varying feeding time on RNAi plates. **(bottom)** Scatter plot showing the relationship between mean oligomer size and total membrane-bound PAR-3 density measured across increasing levels of PAR-3 RNAi depletion. Dotted curve shows a fit of the data to model prediction. **(J) (left)** Representative micrograph of a maintenance phase embryo expressing endogenously-tagged mNeonGreen::PAR-3 and depleted of myosin II heavy chain NMY-2 by RNAi. **(right)** Individual trajectories (yellow traces) for all oligomers detected in the first movie frame. **(K)** Estimated rate constants for dissociation rates of PAR-3 oligomers from the membrane as a function of inferred number of subunits n. Error bars represent the SEM. Data were compiled from n = 3 individual embryos. (L) Relationship between effective dissociation rate and normalized PAR-3 density computed from our empirical measurements. Dashed vertical lines indicate normalized densities of anterior and posterior PAR-3. Dashed horizontal lines indicate the corresponding effective dissociation rates, whose ratio gives the contribution of size-dependent oligomer binding avidity to steady state PAR-3 asymmetries.

This simple model makes several key predictions (Figure 3B-D; see Modeling Supplement for details): First, if oligomers dissociate much more slowly than monomers, as we confirm below (Figure 3K), then the steady-state distribution of oligomer sizes should be approximately exponential, given by *A*_*n*_∼ *A*_1_α^*n*–1^, where *A*_*n*_ is the density of oligomers with n subunits, α = *K_P_ A_1_* measures oligomerization strength, and 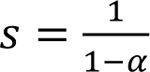 is the mean oligomer size (Figure 3B).

Second, both oligomerization strength α, and mean oligomer size s, should increase with the density of membrane-bound PAR-3 according to:

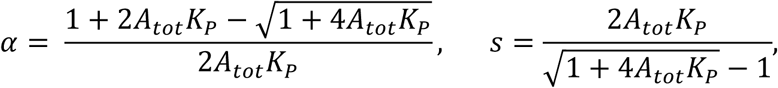

(Figure 3C).

Finally, if the first two predictions hold, and oligomer dissociation rate *k_off_*(*n*) decreases with size, as expected from simple avidity effects ^42,43^, then the effective dissociation rate constant for PAR-3 oligomers, defined as:

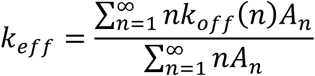

should decrease with increasing PAR-3 density (Figure 3D). This represents a form of positive feedback on PAR-3 accumulation, whose strength is determined by the size-dependence of oligomer dissociation *k_off_*(*n*), and by oligomerization strength through *A*_*n*_ = *A*_1_α^*n*–1^. We will refer to this form of feedback as positive feedback on oligomer binding avidity.

### Intrinsic kinetics of membrane binding and oligomerization produce an approximately exponential distribution of PAR-3 oligomer sizes

To test the first two predictions, we measured the distributions of PAR-3 oligomer sizes on anterior and posterior membranes in *par-1* mutant embryos expressing endogenously-tagged PAR-3::GFP at quasi-steady state during late maintenance phase. We used the intensities of single PAR-3::GFP molecules, measured in the same embryos after photobleaching, as an internal size standard (Figure 3E.F, see Methods).

Consistent with prediction, both the anterior and posterior distributions of oligomer sizes were well fit by single-exponential functions (Figure 3G). The estimated oligomerization strengths (α_*A*_ = 0.730 ± 0.052, α_P_ = 0.424 ± 0.054) and mean oligomer sizes (*S*_*A*_ = 3.48 ± 0.65, *S*_P_ = 1.65 ± 0.16) were higher on the anterior cortex where total PAR-3 density is higher (Figure 3H), and their ratios agreed closely with the predicted dependence of mean oligomer size on PAR-3 densities. We further confirmed this predicted dependence by using RNAi to systematically deplete endogenously tagged PAR-3::GFP, and making paired measurements of total PAR-3 density and mean oligomer size (Figure 3I; see Methods for details). Thus, the exponential distribution of oligomer sizes and the increase in mean oligomer size with PAR-3 density are consistent with a quasi-steady state governed by reversible membrane binding and self-oligomerization of PAR-3.

### Positive feedback on oligomer binding avidity contributes to steady state PAR-3 asymmetry

As described above, positive dependence of mean oligomer size on PAR-3 density, plus negative dependence of oligomer dissociation rate on size, implies positive feedback on PAR-3 accumulation, which we refer to here as positive feedback on oligomer binding avidity. To determine the contribution of this feedback to PAR asymmetries during maintenance phase, we measured the dissociation rate *k_off_*(*n*) as a function of PAR-3 oligomer size. We performed high-speed imaging and particle tracking analysis of PAR-3 endogenously-tagged with mNeonGreen (mNG::PAR-3), using the mean intensities of single molecules to calibrate estimates of oligomer size, as described above (see Methods). To improve the accuracy of particle tracking, we performed these experiments in embryos depleted of the myosin II heavy chain NMY-2 by RNAi to eliminate confounding effects of contractility and cortical flow (Figure 3J; Movie S3). To avoid the confounding effects of photobleaching, we used the intensities of oligomers in the first frame of the imaging sequence to estimate oligomer size n, and the fraction of oligomers in each size class with bound lifetimes > 1 sec to estimate *k_off_*(*n*) (see Methods). Our measurements suggest that *k_off_*(*n*) decreases sharply with oligomer size, becoming negligible for oligomer sizes > 3 subunits (Figure 3K). Using the same data to compare the intensities of PAR-3 oligomers arriving at the membrane with single molecule intensities, we also confirmed that PAR-3 recruitment is dominated by monomer binding (Figure S2A). We achieved comparable results using a strain in which PAR-3 is endogenously-tagged with GFP, although the estimated dissociation rates were slightly higher, likely due to increased particle tracking errors (Figure S2B).

We estimated the contribution of positive feedback on oligomer binding avidity to steady-state PAR-3 asymmetries as follows: At steady state, the local density of PAR-3 is set by the ratio of rate constants for monomer binding and effective dissociation:

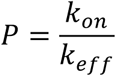

Thus, the ratio of anterior to posterior densities is determined by a product of two factors:

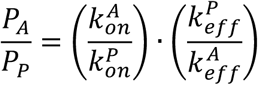

where 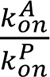, and 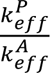 is the ratio of monomer recruitment rates, and α^A^ is the ratio of effective dissociation rates, which measures positive feedback on oligomer binding avidity. Using our measurements of density-dependent oligomerization strengths (α_*A*,_, α_P_; Figure 3H) and size-dependent dissociation (*k_off_*(*n*); Figure 3K) to determine values for *k^A^_eff_* and *k^P^_eff_*, the ratio of effective dissociation rates is 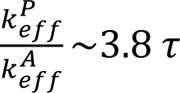. (Figure 3L; see Modeling Supplement for details). Thus, positive feedback on oligomer binding avidity accounts for roughly half of the 8-fold asymmetry in PAR-3 observed in late maintenance phase.

### Kinetics of oligomer unbinding and disassembly set the timescale for approach to steady state during maintenance phase

To further constrain our model, and test its ability to explain the FRAP kinetics and the dynamic approach to steady state observed during maintenance phase, we developed single molecule approaches to distinguish and measure rates of monomer unbinding and oligomer disassembly. Based on previous work and the results above ^17,26,39^, we expect that most posterior PAR-3 molecules are freely-diffusing monomers, whose disappearance reflects the faster kinetics of membrane unbinding. In contrast, we expect that most anterior PAR-3 molecules are associated with stably-bound slow-moving PAR-3 oligomers, and their disappearance reflects the slow kinetics of oligomer disassembly. To confirm this, we imaged single molecules of endogenously-tagged PAR-3::GFP using high laser power at 50 msec intervals to ensure accurate particle tracking (Figure 4A;Movie S4; see Methods). To quantify disappearance, we constructed release curves by plotting the numbers of molecules with lifetimes > τ as a function of τ (Figure 4B).

**Figure 4.**
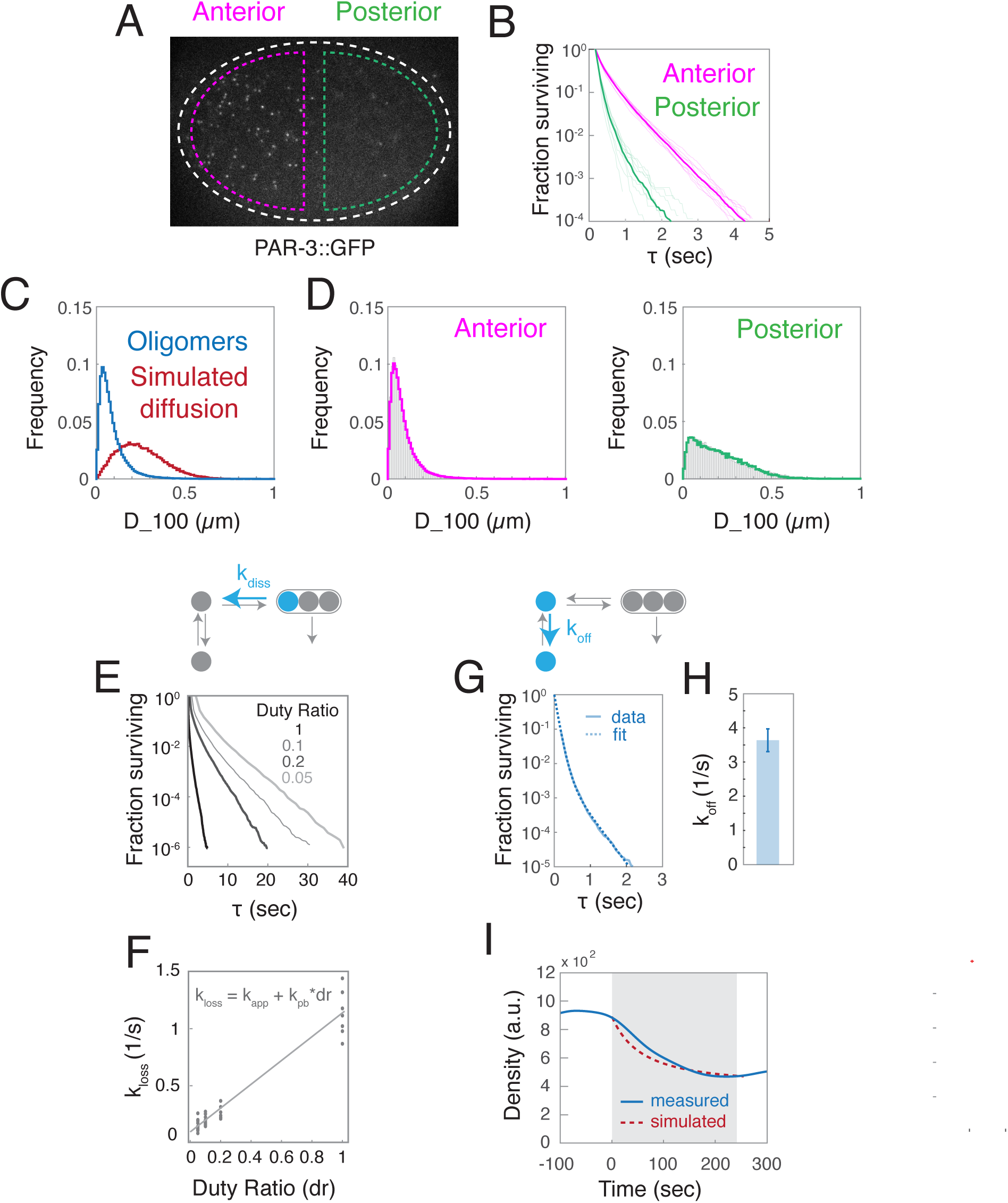
Membrane unbinding and depolymerization set the timescale for slow approach to steady state distributions of PAR-3 during maintenance phase **(A)** Representative image showing single molecules of PAR-3::GFP imaged at high laser power and short exposure times to ensure accurate tracking of fast diffusing molecules. **(B)** Release curves measured for single molecules of PAR-3::GFP on the anterior and posterior cortex of wild type embryos under continuous illumination. Thin traces show release curves for individual embryos (n = 8); Thick traces show data pooled for all embryos. Data were normalized by total numbers of anterior tracks. **(C)** Distributions of displacements after 100 msec (D_100) for (blue) tracked PAR-3 oligomers (n = 7 embryos) and (red) simulated Brownian diffusion with diffusivity D = 0.1 µm^2^/sec. **(D)** D_100 distributions for (left) anterior and (right) posterior molecules pooled from 8 embryos. Solid curves show least squares best fit to weighted distributions for oligomers and for simulated diffusion. **(E)** Release curves for PAR-3::GFP molecules imaged with fixed laser power and exposure times but different duty ratios. Each curve represents data pooled from multiple embryos: (dr = 1, n = 7; dr = 0.2; n = 7, dr = 0.1; n = 12; dr = 0.05, n = 13). **(F)** Loss rate constants (*k_loss_*) measured for different duty ratios by fitting the linear portions of the release curves (see Methods for details). The dashed line represents a linear least-squares fit of the data to the equation *k_loss_* = *k_app_* + *k_pb_* ∗ *dr*. **(G)** Release curves measured for posterior single molecules of endogenously-tagged PAR-3::GFP in control embryos (n = 9). Dashed lines indicate data fit to a weighted sum of two exponentials. **(H)** Corresponding estimate of effective *k_off_* from the data in (G). Error bars indicate 95% confidence intervals obtained from bootstrap sampling. **(I)** Comparison of the approach to steady state measured in *par-1* mutant embryos (blue curve) and the simulated approach (dashed red curve) given measured values for *k_off_*(*n*) and *k_diss_*.

To quantify short-term mobilities, we measured distributions of single-molecule displacements after 100 msec (D_100), and fit these distributions to weighted sums of D_100 distributions measured for PAR-3 oligomers (Figure 4C, blue curve), or computed for simulated Brownian diffusion (Figure 4C, red curve).

Consistent with our expectations, the mobilities of anterior PAR-3 molecules closely resemble those measured for PAR-3 oligomers (Figure 4D, left), while posterior mobility distributions reflect a dominant contribution (∼ 75%) from Brownian diffusion at ∼0.1 µm^2^/sec (Figure 4D, right). Posterior release kinetics are much faster than anterior release kinetics (Figure 4B), confirming that membrane unbinding is faster than oligomer disassembly, although photobleaching obscures the magnitude of this difference. Plotting release curves on a logarithmic scale reveals a single dominant release rate for anterior molecules with lifetimes > 1 second, reflecting the slow kinetics of oligomer disassembly. In contrast, posterior molecules release with a mix of fast and slower kinetics (Figure 4G), both of which likely reflect monomer dissociation, because oligomerization-defective PAR-3 molecules release with a similar kinetics (Figure S3D and below). Importantly, these results were relatively insensitive to the intensity threshold used to detect single molecules (Figure S3).

Based on these observations, we used the following strategy to estimate rate constants for oligomer disassembly (*k_diss_*) and monomer unbinding (*k_off_*) from the observed release kinetics. First, we imaged anterior PAR-3 molecules, holding laser power and exposure time constant while varying the duty ratio of exposure (*dr*). For each duty ratio, the resulting release curves were well-fit by single exponentials for lifetimes > 1 sec, yielding an estimated rate constant *k^a^_loss_* for the observed loss of anterior PAR-3 molecules (Figure 4E). The observed loss rate is a sum of two contributions: *k^a^_loss_* = *k_app_* + *k_pb_* ∗ *dr*, where *k_pb_* ∗ *dr* accounts for photobleaching, and *k_app_*describes the apparent rate of a two-step process in which PAR-3 monomers first dissociate from oligomers and then unbind the membrane. We extracted estimates of *k_pb_* (1.04 ± 0.04 sec^-1^) and *k_app_* (0.08 ± 0.02) from the slope and y-intercept respectively of linear fits to our estimates of *k^a^_loss_ v dr* (Figure 4F; see Methods for details).

We fit posterior release curves obtained in streaming mode (dr = 1) to a sum of exponentials to estimate an effective loss rate for posterior monomers *k^p^_loss_*, and then subtracted the measured photobleaching rate *k_pb_* to estimate an effective monomer unbinding rate *k_off_* = *k^p^_loss_* − *k_pb_* = 3.75 ± 0.34 *sec*^-1^ (Figure 4G; see Methods for details), which is comparable to the value measured above by tracking PAR-3 clusters (k_off_(1) = 2.7/sec; Figure 3K). Since k_off_ is > 30-fold slower than k_app_, oligomer unbinding must be rate-limiting for dissociation of anterior PAR-3 molecules, and therefore *k_diss_*∼*k_app_* = 0.08/*sec*.

Using the measured values for *k_off_*(*n*), *k_diss_* and α to constrain a steady-state flux balance analysis, we inferred that no more than 17% of PAR-3 monomers bind directly to existing oligomers, although the actual value is likely significantly lower, justifying the choice to neglect direct binding in our model. (see Modeling Supplement, Section 1.1.2). We also compared these single measurements to the measured kinetics of FRAP recovery within PAR-3 oligomers (Figure 2I-K). If monomers dissociate very rapidly, and oligomers very slowly, as our measurements show, then on intermediate timescales (10s of seconds) the fluorescence recovery should be dominated by the kinetics of oligomer disassembly. Indeed, the predicted half time for recovery of bleached fluorescence within oligomers is 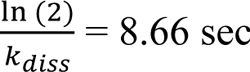, which agrees well with the measured half time for FRAP recovery. Furthermore, if PAR-3 assembles linear oligomers, then only terminal subunits can undergo rapid exchange. Given the measured oligomer size distribution, 43% of anterior oligomer subunits are terminal, which agrees well with the ∼50% recovery observed in our FRAP experiments (see Modeling Supplement). Thus, dynamic exchange of terminal subunits on linear oligomers could explain both the kinetics and the extent of the observed recovery.

Finally, we simulated the dynamic approach to steady state during maintenance phase, assuming a constant rate of monomer binding, and using the measured values for *k_off_*(*n*), *k_diss_* and α. We observed good agreement between simulated and measured dynamics (Figure 4I), confirming that the intrinsic kinetics of PAR-3 membrane binding and oligomerization are sufficient to explain the timescale of approach to steady state during maintenance.

### Asymmetric monomer recruitment contributes to PAR-3 asymmetries during maintenance phase

The results above show that positive feedback on oligomer binding avidity accounts for roughly half the PAR-3 asymmetries observed *par-1* mutant embryos. Theoretical analysis shows that, regardless of strength, this form of feedback cannot stabilize an asymmetric distribution of oligomers, although it can reduce the amount additional feedback required to do so ^44^. Thus, additional forms of feedback are required to explain the observed asymmetries.

Membrane-bound PAR-3 could feed back locally (through one or more molecular intermediaries) to promote monomer binding and/or inhibit monomer dissociation. To test these possibilities, we used single molecule imaging to measure relative rates of PAR-3 monomer binding and dissociation on anterior and posterior membranes during maintenance phase. To avoid the potentially confounding effects of oligomerization, we performed these experiments in a transgenic strain expressing a variant of PAR-3 (PAR-3<u>^V80D,D138K^</u>::GFP) containing two point mutations in the conserved N-terminal CR1 domain that disrupt homo-oligomerization of PAR-3^34,35,37^. (Figure 5A,B; Movie S5). This allowed us to quantify binding and dissociation of monomeric PAR-3<u>^V80D,D138K^</u>::GFP in the context of endogenously-expressed PAR-3 and normal PAR asymmetries ^37^.

**Figure 5.**
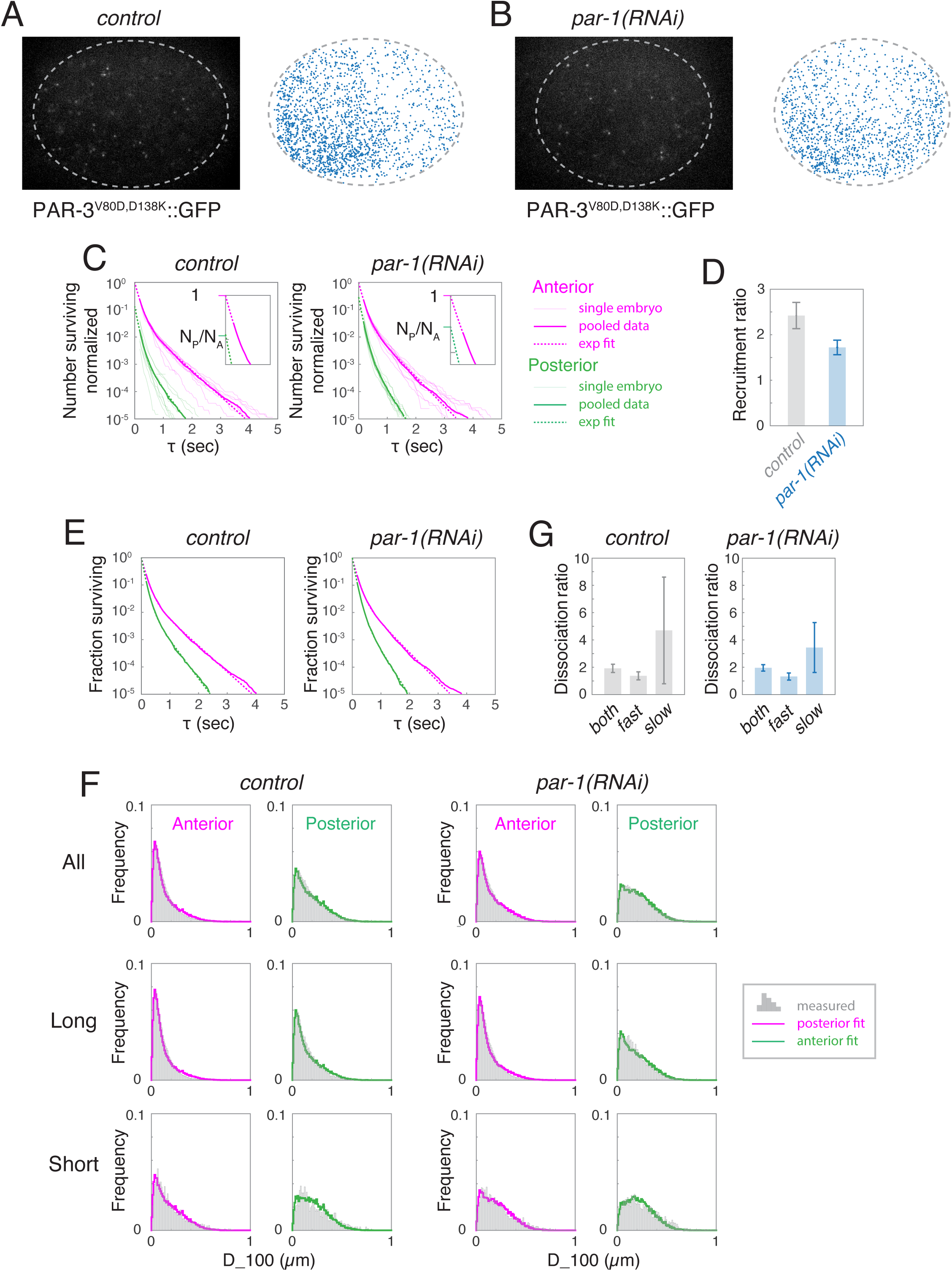
Asymmetric recruitment of PAR-3 monomers underlies maintenance of PAR-3 asymmetries. **(A,B)** Representative images and distribution of single molecule appearance events for oligomerization-defective PAR-3^V80D,D138K^ in **(A)** control and **(B)** *par-1(RNAi)* embryos, detected by single molecule imaging. Each blue dot represents a single binding event. **(C)** Release curves for control (n = 9) and *par-1(RNAi)* (n = 10) embryos. Faint lines show data for individual embryos, thick lines for pooled data, and dashed lines indicate the exponential fits to pooled data. Inset shows the extrapolated zero-crossings from which the total numbers of binding events were inferred. Anterior and posterior data were normalized to total numbers of anterior binding events to highlight the recruitment asymmetry. **(D)** A:P recruitment ratios measured from the data in **(C)**. Error bars indicate 95% confidence intervals obtained from bootstrap sampling. **(E)** Pooled release curves and exponential fits with anterior and posterior data separately normalized by total number of observed trajectories to highlight differences in release rates. **(F)** Distributions of short-term displacements (D_100) for anterior and posterior molecules in control and *par-1(RNAi)* embryos, measured over all trajectories (top row), long trajectories (middle row; duration > 2 sec), and short trajectories (bottom row; duration = 250 msec). Solid curves show data fits to weighted sums of D_100 distributions for oligomers and for simulated Brownian diffusion with diffusivity D = 0.1 µm<u>^2^</u>/sec (as in Fig 4C). **(G)** A:P ratios of effective monomer dissociation rates for fast and slow components (both) and for fast and slow components alone. Error bars indicate 95% confidence intervals obtained from bootstrap sampling.

To measure recruitment rates, we performed single molecule imaging and particle tracking analysis as described above, and counted the total number of tracked molecules appearing on the anterior or posterior membrane per unit area and time. To avoid counting cytoplasmic molecules that diffuse transiently into the focal plane without binding, we only considered molecules with tracked lifetimes > 200 msec, and extrapolated from exponential fits to the release curves to estimate the total number of binding events (see Figure 5C insets, Methods for details).

With this approach, we measured an approximately 1.8-fold A:P difference in PAR-A^3V80D,D138K::GFP^ recruitment rates in *par-1(RNAi)* embryos 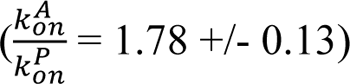, and slightly higher ratios in control embryos (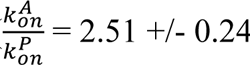; Figure 5D), consistent with the greater PAR-3 asymmetries observed in controls. We obtained similar results using a PAR-3 transgene containing a deletion of 14 amino acids within the CR1 domain that also abolishes self-oligomerization (Figure S4). These data suggest that monomeric PAR-3 binds the membrane at higher rates where endogenous PAR-3 is already enriched, consistent with positive feedback on PAR-3 recruitment. Moreover, taken together, the A:P ratios in effective dissociation rate (3.8) and recruitment rates (∼1.8) are nearly sufficient to account for the total PAR-3 asymmetry observed at late maintenance phase in *par-1* mutant embryos.

### Asymmetric monomer dissociation contributes to maintaining PAR-3 asymmetries

Comparing anterior and posterior release curves normalized by total numbers of observed molecules (Figure 5E) revealed that anterior PAR-3<u>^V80D,D138K^</u>::GFP molecules dissociate with slower overall kinetics than posterior molecules. For trajectories with lifetimes > 200 msec, anterior and posterior release curves were well approximated by a weighted sum of two exponential functions (dashed lines in Figure 5E), revealing the existence of at least two distinct membrane binding states with different release kinetics. Similarly, the distributions of short-term displacements (D_100) were reasonably well fit by a weighted sum of two distributions: one representing simulated Brownian diffusion with D = 0.1 µm^2^/sec, and the other representing the sub-diffusive mobility of cortically associated factors, such as PAR-3 oligomers (Figure 5F, top row). Finally, longer trajectories (>2 sec) were enriched for the sub-diffusive mobility state, while shorter trajectories (< 250 msec) were enriched for the diffusive mobility state (Figure 5F, middle and bottom rows). Together, these data suggest that PAR-3<u>^V80D,D138K^</u>::GFP monomers occupy two distinct membrane-bound states: a diffusive state with fast release kinetics and, unexpectedly, a sub-diffusive state with slower release kinetics.

From the exponential fits for *par-1(RNAi)* embryos shown in Figure 5E, we estimate a ∼ 2-fold asymmetry in effective release rates for all PAR-3^V80D,D138K^::GFP monomers (Figure 5G, both). However, most of this difference is due to the slow-releasing molecules (compare Figure 5G, fast and slow). If all monomers bind PAR-3 oligomers with the same kinetics, then the combination of asymmetric recruitment and release will produce an asymmetry in PAR-3 density that is 1.5 fold higher than the value measured in *par-1* mutant embryos. However, sub-diffusive PAR-3 monomers are expected to encounter and bind PAR-3 oligomers at far lower rates than fast-diffusing monomers, and should therefore make much smaller contributions to oligomer assembly. Indeed, the mean lifetime of the slow dissociating sub-diffusive pool is ∼ 3 sec.

Considering only those molecules that remain bound for at least this long, the mean displacement in 3 sec is ∼ 0.6µm, which is a little more than half the mean distance between oligomers, implying that the majority of these molecules dissociate before they bind another PAR-3 molecule. If only the fast-releasing pool of PAR-3 monomers contributes to oligomer assembly, the A:P release ratio is ∼1.3 (Figure 5G, fast), and the steady-state asymmetry is predicted to be ∼8, which matches the measured value. In summary, our data suggest that asymmetric dissociation of PAR-3 monomers contributes to sustaining PAR-3 asymmetries during maintenance phase, but the magnitude of this contribution is likely to be fairly small.

### The combination of positive feedback on PAR-3 monomer recruitment and oligomer binding avidity are necessary and sufficient to produce stable PAR-3 asymmetries

To determine whether the combination of positive feedback on monomer recruitment and positive feedback on oligomer binding avidity are sufficient to explain the dynamically stable PAR-3 asymmetries observed in *C. elegans*, we considered a spatially distributed version of the model in Figure 3A, and added a simple form of feedback (cyan arrow in Figure 6A), in which the rate constant for monomer recruitment is given by:

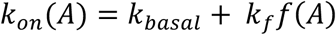

**Figure 6.**
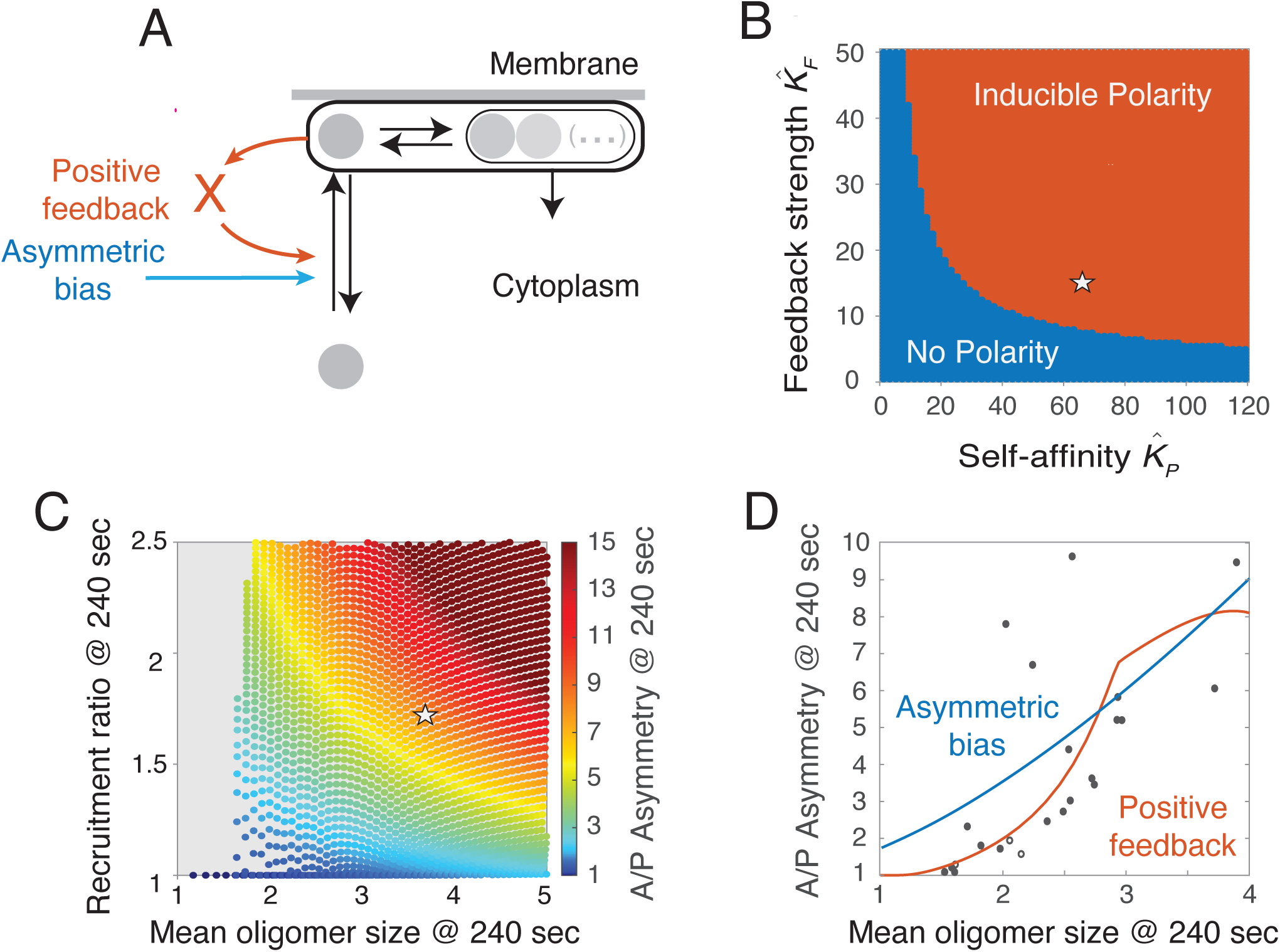
Positive feedback on PAR-3 monomer recruitment and size-dependent dissociation of oligomers, are both necessary and sufficient to stabilize PAR-3 asymmetries. **(A)** Modified versions of the reaction diagram in Figure 3A, showing two alternative assumptions for the mechanism that underlies asymmetric recruitment: (blue arrow): Membrane-bound PAR-3 feeds back (likely through intermediaries X) to promote recruitment of PAR-3 monomers to the membrane. (orange arrow): A fixed asymmetric input, whose strength is independent of PAR-3 densities, biases recruitment to the anterior cortex. **(B)** Theoretical phase plane derived for the version of the model with positive feedback on monomer recruitment, showing how the ability to polarize depends on K^^^_F_ and K^^^_P_, which tune the strengths of feedback on monomer recruitment and oligomer binding avidity respectively. No polarity: Only the spatially uniform steady state is stable. Inducible polarity: Spatially uniform and polarized states are both stable, and local depletion of PAR-3 can induce a transition from the spatially uniform to the polarized state (see text for details). White star indicates the values that reproduce the experimentally measured mean oligomer size and recruitment asymmetry in the *C. elegans* zygote. **(C)** Experimental phase plane computed by simulating PAR dynamics for 240 sec during maintenance phase. Colored dots indicate the predicted asymmetry as a function of monomer recruitment asymmetry and mean oligomer size measured at the 240 sec timepoint (see text for details). White star indicates the prediction for experimentally measured mean oligomer size and recruitment asymmetry in the *C. elegans* zygote. **(D)** Comparison between predicted and measured relationships between mean oligomer size and PAR-3 asymmetry in late maintenance phase. Blue and orange curves show the predicted asymmetries, after 240 sec of simulated time, as a function of total PAR density, for the positive feedback (blue curve) and fixed asymmetric bias (orange curve) scenarios shown in Figure 6A. Black circles show mean oligomer size and PAR-3 asymmetry measured for individual embryos in late maintenance phase. The two unfilled circles indicate the only two embryos in which PAR-3 was not visibly polarized in early maintenance phase.

Where *k_basal_* is the rate constant for basal (PAR-3 independent) recruitment, and *k_f_* tunes the strength of positive feedback. Casting the model into dimensionless form (see Modelling Supplement), this becomes:

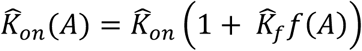

where K^^^_on_ is the dimensionless rate constant for basal recruitment, and K^^^_f_ the dimensionless feedback strength. Because recruitment asymmetries of PAR-3 are similar in early and late maintenance phase, despite a nearly two-fold change in the anterior PAR-3 density (Figure S5A), we assumed a simple form of saturating feedback governed by:

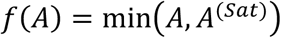

where *A*^(sat)^ is the density of PAR-3 above which feedback saturates.

Using local perturbation analysis (LPA^45^), we found that only two types of dynamical behavior are possible: Either the spatially uniform steady state is the only stable state (***No polarity***), or spatially uniform and asymmetric stable states coexist, and transient local depletion of PAR-3 can induce a transition from the spatially uniform to the polarized state (***Inducible polarity***). Importantly, inducible polarity corresponds to the behavior observed in *C. elegans* zygotes, in which spatially uniform distributions of PAR-3 are stable in the absence of a functional sperm cue, but depletion of posterior PAR-3 by cortical flow induces the transition to a spatially asymmetric distribution that remains stable after the cue is gone, even in the absence of PAR-1.

To determine how the two forms of feedback shape these outcomes, we analyzed the predicted steady state behavior as a function of K^^^_F_, which quantifies the strength of feedback on monomer recruitment, and K^^^_P_, a dimensionless measure of PAR-3 self-affinity that quantifies the strength of feedback on oligomer binding avidity. We set *A*^(sat)^ so that positive feedback on recruitment saturates at 80% of the steady state PAR-3 density. For each value of K^^^_F_and K^^^_P_, we set K^^^_on_so that 10% of total PAR-3 is enriched on the membrane at the spatially uniform steady state, and then we used LPA to determine the predicted dynamics. The resulting phase diagram reveals that the two forms of feedback act synergistically - increasing the strength of one form of feedback reduces the need for the other – but that both are absolutely required for inducible polarity (Figure 6B).

Because PAR-3 asymmetries approach, but do not reach, steady state during maintenance phase, we cannot directly compare our experimental measurements to the phase diagram shown in Figure 6B. For a more direct comparison, we assigned values to K^^^_on_and *A*^(sat)^as above, and we used the empirically measured rate constants for oligomer unbinding K^(*n*)^_off_ and disassembly K_diss_. For each combination of feedback strengths K^^^_F_and K^^^_P_in Figure 6B, we initialized simulations using the distribution of PAR-3 measured at the beginning of maintenance phase. We simulated the approach to steady state for 240 seconds. Then, from the final state, we extracted the predicted recruitment ratio, mean oligomer size, and A/P asymmetry, and plotted the predicted dependence of A:/P asymmetry on recruitment ratio and mean oligomer size, all measured after 240 sec of simulated time (Figure 6C). Strikingly, given the recruitment ratio (1.8) and mean oligomer size (3.7) measured in late maintenance, the model predicts an A:P asymmetry of ∼8 (magenta disk in Figure 6C), which is close to the measured value. Furthermore, the values of K^^^_F_and K^^^_P_ for which simulations reproduce the measured recruitment ratio and mean oligomer size (14.5 and 62 respectively) locate the embryo well within the inducible polarity regime (magenta disk in Figure 6B). Importantly, these results were relatively insensitive to the exact choice of the feedback saturation point *A*^(sat)^. We conclude that if positive feedback drives the asymmetric recruitment of PAR-3, then the combined strengths of positive feedback on monomer recruitment and on oligomer binding avidity are sufficient to support the induction of dynamically stable unipolar PAR-3 asymmetries in response to a transient local depletion of PAR-3, in the absence of posterior inhibition by PAR-1.

In principle, the asymmetric recruitment of oligomerization-defective PAR-3 could reflect a fixed bias independent of local PAR-3 density, rather than positive feedback (compare blue and red arrows in Figure 6A). To distinguish these possibilities, we performed a mathematically controlled comparison ^46^ between our model and one in which positive feedback on monomer recruitment is replaced by a fixed recruitment bias that reproduces an identical wild-type solution – i.e., one that reproduces same recruitment asymmetry, mean oligomer size and PAR-3 asymmetry after 240 seconds of simulated time from the same initial conditions. Specifically, we compared the predicted response of these two models to systematic variation in mean oligomer size, produced by depletion of total PAR-3 levels (Figure 6D). For the positive feedback scenario, the model predicts that as mean oligomer size falls below a threshold level, the stably asymmetric steady state disappears, and PAR-3 asymmetries approach a minimal value of 1 (they do not reach the minimum value because of the finite simulation time). In contrast, for a fixed recruitment bias, the model predicts that PAR-3 asymmetry approaches a constant minimum value of ∼ 1.8 as the mean oligomer size approaches 0.

To distinguish these possibilities, we used RNAi to systematically vary total PAR-3 levels (and thus mean oligomer size; see Figure 3G) in *par-1* mutant embryos expressing endogenously tagged PAR-3::GFP. Importantly, depletion of PAR-3 prevents cortical flows during polarity establishment in otherwise wild-type embryos ^11^, but not in *par-1* mutant embryos (Figure S5B), allowing us to analyze PAR-3-depleted embryos that segregate residual PAR-3 into an anterior cap during polarity establishment phase (Figure S5C). For each sampled embryo, we visually confirmed a polarized distribution of PAR-3 in early maintenance phase and then made paired measurements of the mean oligomer size and the PAR-3 asymmetry in late maintenance phase (see Methods for details). The measured relationship between PAR-3 asymmetry and mean oligomer size closely matched the response predicted for the positive feedback scenario, both qualitatively and quantitatively (Figure 6D). In particular, PAR-3 asymmetries approach the limiting value of 1 as mean oligomer size decreases, which is completely inconsistent with a fixed recruitment asymmetry. Thus, regardless of the underlying mechanism, these results strongly support a model in which positive feedback on monomer binding drives asymmetric recruitment of PAR-3, and they show that, when combined with positive feedback on membrane binding avidity, this feedback is sufficiently strong to sustain a dynamically stable unipolar state in embryos lacking posterior inhibition by PAR-1.

### PAR-6/PKC-3 are required for positive feedback on PAR-3 recruitment and stable PAR-3 asymmetries

Because positive feedback on PAR-3 recruitment does not require an intact CR1 domain, it likely involves the interaction of PAR-3 with additional factors. The other anterior polarity proteins PAR-6, PKC-3 and CDC-42 are attractive candidates. PAR-6 and PKC-3 form an obligate heterodimer (PAR-6/PKC-3) that can bind directly to either PAR-3 or CDC-42 ^47–54^. Both PAR-3 and CDC-42 ^17,55^ are required to recruit PAR-6/PKC-3 into an active complex with CDC-42 during maintenance phase, and the PAR-6/PKC-3 heterodimer is thought to cycle between PAR-3-bound and CDC-42-bound states during polarization ^16,39,55,56^. suggesting that PAR-6/PKC-3, and/or CDC-42, could act as intermediaries to promote positive feedback on PAR-3 recruitment.

To test these possibilities, we first examined the effects of depleting PAR-6 on PAR-3 asymmetries in *par-1* mutant and heterozygous control embryos (Figure 7). Consistent with previous reports ^11,16,57^, PAR-3 oligomers accumulated on the membrane during polarity establishment phase in control (*par-1/+*) embryos depleted of PAR-6, although cortical flows and anterior enrichment of PAR-3 were significantly reduced (Figure 7A; *par-6(RNAi)*, Movie S6). However, these asymmetries were completely lost during maintenance phase. This loss of asymmetry was accompanied by both a rapid decrease in PAR-3 density and oligomer size on the anterior cortex, and by the rapid transport of PAR-3 oligomers towards the posterior pole by aberrant posterior-directed cortical flows, which depend on activation of myosin II by the kinase MRCK-1^17,22^ (Figure 7A, Movie S6). However, co-depleting PAR-6 and MRCK-1 produced a similarly rapid decrease in anterior oligomer size and a complete loss of PAR-3 asymmetry during maintenance phase, even in the absence of aberrant cortical flow (Figure 7A,B; *par-6(RNAi);mrck-1(RNAi)*, Movie S6).

**Figure 7.**
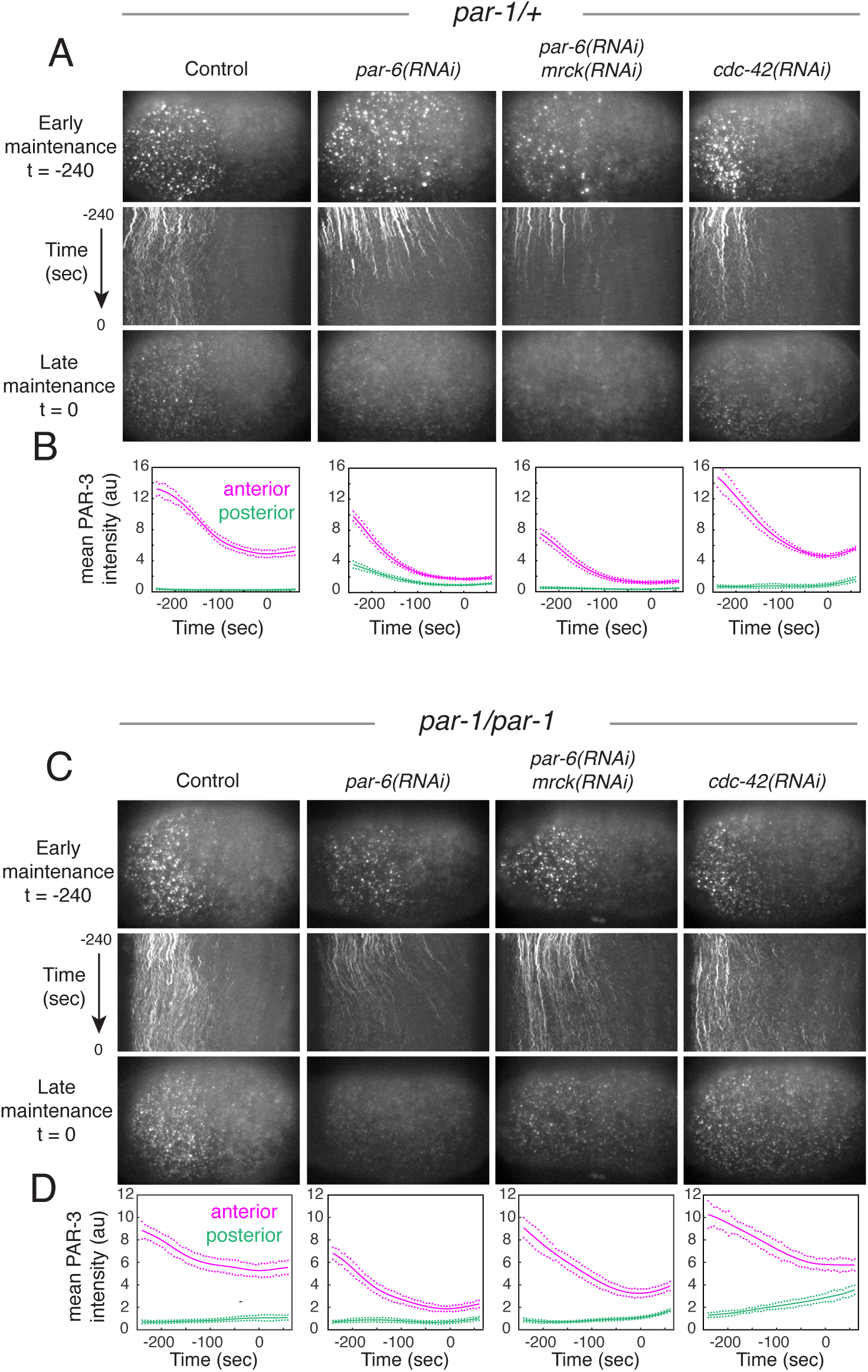
PAR-6/PKC-3 and CDC-42 are required to maintain PAR-3 asymmetries during maintenance phase. **(A)** Representative micrographs showing PAR-3 oligomer sizes and distribution during maintenance phase in *par-1* heterozygote (*par-1/+*) embryos for the indicated RNAi depletions. Top and bottom micrographs show the distribution of PAR-3 oligomers in early (−240 sec) and late (0 sec) maintenance phase. Time is measured relative to the point where anterior density of PAR-3 reached a minimum in late maintenance phase (see Methods for details). Kymographs show how the distribution changes over the 240 seconds between these time points. Diagonal streaks in the kymographs indicate transport of PAR-3 oligomers by cortical flow. **(B)** Corresponding plots of total fluorescence density of PAR-3 within segmented speckles during maintenance phase. Bold lines indicate the mean; green and magenta dots indicate the standard error of the mean across individual embryos (control: n = 8, *cdc-42(RNAi)*: n = 7, *par-6(RNAi):* n = 6, *par-6(RNAi)*; *mrck-1(RNAi*): n = 8). **(C)** Representative micrographs showing PAR-3 oligomer sizes and distribution during maintenance phase in *par-1* homozygous (*par-1/par-1*) embryos for the indicated RNAi depletions. Data are presented as described in **(A)**. **(D)** Corresponding plots of total fluorescence density of PAR-3 within segmented speckles during maintenance phase. Bold lines indicate the mean; green and magenta dots indicate the standard error of the mean across individual embryos (control: n = 10, *cdc-42(RNAi)*: n = 6, *par-6(RNAi):* n = 9, *par-6(RNAi)*; *mrck-1(RNAi*): n = 14).

The decrease in PAR-3 density and oligomer sizes produced by depleting PAR-6 in *par-1* heterozygotes could be due to increased inhibition by PAR-1^12^, which accumulates uniformly in embryos lacking PAR-6/PKC-3^58^. However, depleting PAR-6 in *par-1* homozygous mutant embryos produced a similarly rapid and complete loss of PAR-3 oligomers during maintenance phase (Figure 7C,D; *par-6(RNAi),* Movie S7). We also observed a similarly rapid loss of PAR-3 asymmetry in *par-1* homozygotes co-depleted of PAR-6 and MRCK-1, although the loss was incomplete at late maintenance phase (Figure 7C,D; *par-6(RNAi);mrck-1(RNAi)*, Movie S7)).

Thus, PAR-6/PKC-3 play a central role in maintaining PAR-3 asymmetries during maintenance phase independent of cortical flow and posterior inhibition by PAR-1.

PAR-6/PKC-3 could maintain PAR-3 asymmetries by inhibiting oligomer disassembly and/or monomer dissociation, and/or by promoting monomer recruitment. Using the single molecule approaches described above, we found there was no significant difference in oligomer disassembly rates (*k_diss_*) during early maintenance phase in PAR-6-depleted embryos compared to untreated controls (Figure 8A,B). In contrast, the asymmetric recruitment of PAR-3<u>^V80D,D138K^</u>::GFP during early maintenance phase was abolished in embryos co-depleted of PAR-6 and MRCK-1 (Figure 8C), even though endogenous PAR-3 is highly enriched on the anterior cortex during early maintenance phase in these embryos (Figure 7A,B). There was also a decrease in ratio of release rates for PAR-3<u>^V80D,D138K^</u>::GFP (Figure S6A-D). We conclude that PAR-6/PKC-3 stabilize PAR-3 asymmetries during maintenance phase primarily by promoting positive feedback on PAR-3 monomer recruitment.

**Figure 8.**
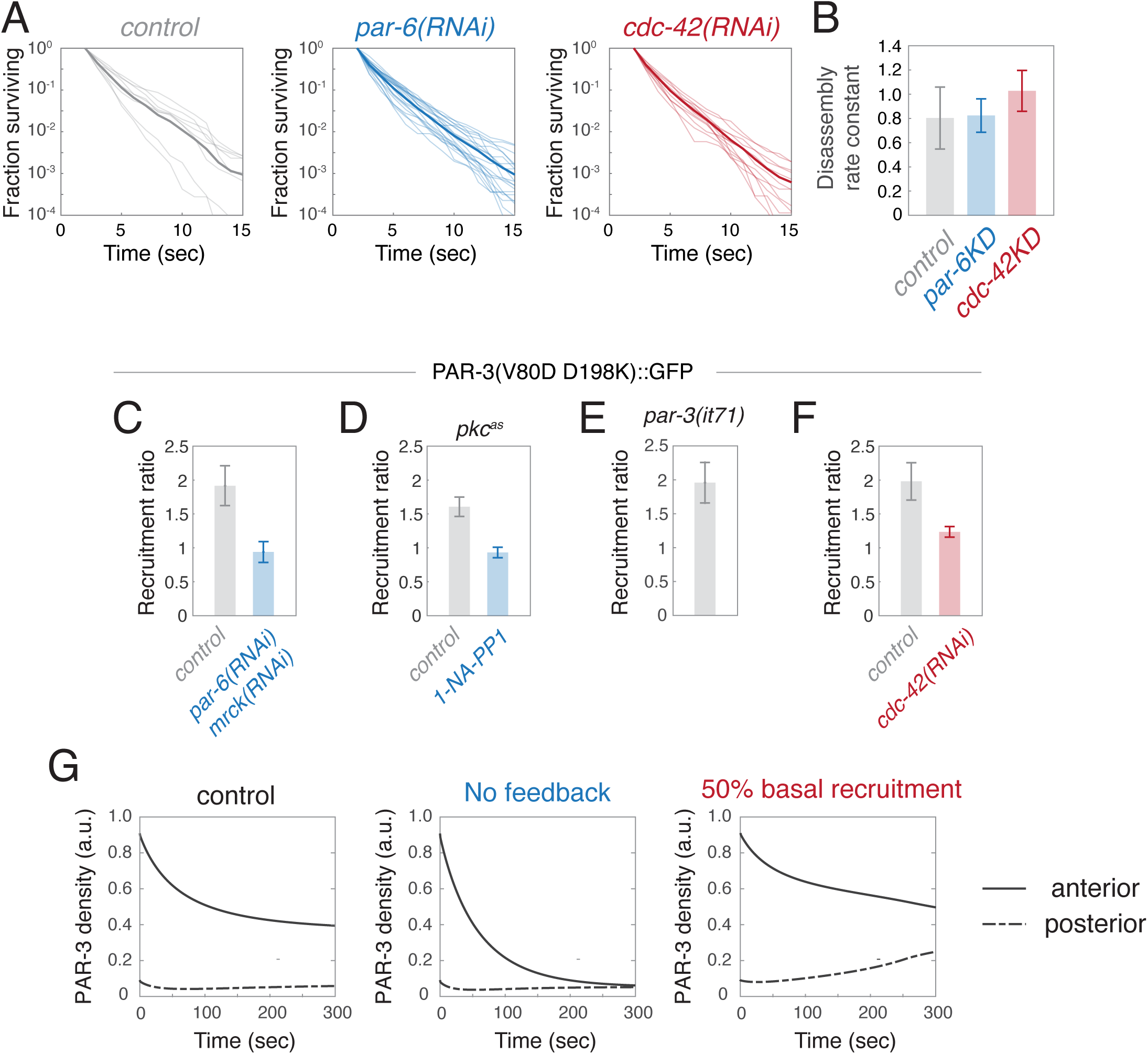
PAR-6/PKC-3 and CDC-42 stabilize PAR-3 asymmetries by promoting asymmetric recruitment and dissociation of PAR-3 monomers. **(A)** Release curves measured for single molecules of transgenic PAR-3::GFP measured in control, *par-6(RNAi)* and *cdc-42(RNAi)* embryos during maintenance phase under conditions that reveal slow-release kinetics of oligomer-bound molecules. All data were collected under identical conditions. Thin curves show data from single embryos. Thick curves show pooled data: (control: n = 8, *cdc-42(RNAi)*: n = 12, *par-6(RNAi)*: n = 14). **(B)** Estimates of oligomer disassembly rate constants (*k_diss_*) from the data shown in **(A)** after correction for photobleaching (see Methods). Error bars represent 95% confidence intervals obtained by bootstrap sampling. **(C-F)** A:P recruitment ratios measured for oligomerization-defective PAR-3::GFP during late maintenance phase under different experimental conditions. Error bars represent 95% confidence intervals obtained by bootstrap sampling. **(C)** Control (n = 9) and *par-6(RNAi);mrck-1(RNAi)* (n = 12). **(D)** Mutant embryos endogenously expressing an analogue-sensitive form of PKC-3 and transgenically expressing oligomerization-defective PAR-3::GFP, and treated with DMSO (control; n = 8) or 50µM ATP analog (1-NA-PP1; n = 6). **(E)** *par-3* mutant embryos transgenically expressing oligomerization-defective PAR-3::GFP (n = 6). **(F)**. Control (n = 6) and *cdc-42(RNAi)* (n = 6). **(G)**. Simulations showing predicted changes in anterior and posterior PAR-3 densities during maintenance phase starting from the initial distributions observed at the onset of maintenance phase. Left panel: Control conditions; Middle panel: Feedback strength set to zero to mimic *par-6(RNAi)*; *mrck-1(RNAi*); Right panel: basal recruitment rate adjusted to mimic *cdc-42(RNAi)*.

To test this idea further, we set the feedback strength K^^^_f_= 0 in our empirically constrained model to mimic the complete loss of feedback on monomer recruitment, and simulated PAR-3 dynamics from the initial distribution measured at late establishment phase in *par-1* homozygote embryos co-depleted of PAR-6 and MRCK (Figure 7C; *par-6(RNAi);mrck-1(RNAi)*). As expected from the phase diagrams in Figure 6B&C, the model predicts the complete loss of asymmetry, but the time course is too slow to reach completion during maintenance phase (Figure 8G), consistent with the incomplete loss of asymmetry observed at late maintenance phase in these embryos (Figure 7D). These results further support the conclusion that PAR-6/PKC-3 act as a key intermediary to support positive feedback of PAR-3 on its own recruitment and to promote dynamically stable PAR-3 asymmetries.

To further probe the mechanism underlying this regulation, we examined PAR-3<u>^V80D,D138K^</u>::GFP recruitment and release in embryos endogenously expressing an ATP analogue-sensitive mutant of *pkc-3* (*pkc-3^as^*^16^). Asymmetric recruitment of PAR-3<u>^V80D,D138K^</u>::GFP was slightly reduced, relative to the wild type, in permeabilized *pkc-3^as^* embryos treated with a vehicle control, but it was completely abolished by treatment of permeabilized embryos with ATP analogue 1-NA-PP1 (Figure 8D). In contrast, the same treatment had no effect on monomer release ratios (Figure S6E-H). We conclude that positive feedback on monomer recruitment requires PKC-3 activity.

Previous studies have shown that a fraction of embryos expressing only an oligomerization-defective form of PAR-3 can divide asymmetrically and complete development^37,39^. In embryos endogenously expressing a point-mutated oligomerization-defective form of PAR-3, a subset of embryos exhibits polarized PAR-6/PKC-3, suggesting that oligomerization-defective PAR-3 is sufficient to support minimal PAR asymmetries ^39^. To test whether oligomerization-defective PAR-3 can support positive feedback on monomer recruitment, we measured asymmetric recruitment of PAR-3<u>^V80D,D138K^</u>::GFP, expressed in homozygous *par-3(it71)* mutant zygotes ^37^.

For *par-3(it71);* PAR-3<u>^V80D,D138K^</u>::GFP zygotes that divided asymmetrically, the recruitment asymmetry measured in late maintenance phase was comparable to that measured in controls (Figure 8E), suggesting that monomeric PAR-3 is sufficient for positive feedback on its own recruitment.

### CDC-42 promotes dynamically stable PAR-3 asymmetries by tuning basal rates of PAR-3 recruitment

We also assessed the consequences of depleting CDC-42 in *par-1* mutant and heterozygous control embryos. In *par-1* heterozygotes depleted of CDC-42, PAR-3::GFP asymmetries formed during polarity establishment phase and relaxed toward lower levels during maintenance phase, as observed in controls, although overall density was slightly higher than in controls (Figure 7A,B; Movie S6; compare control and *cdc-42(RNAi)*). PAR-3::GFP was also enriched during polarity establishment in *par-1* homozygotes depleted of CDC-42, but these asymmetries were almost completely lost during polarity maintenance phase (Figure 7C,D; *cdc-42(RNAi)*, Movie S7). Unlike in PAR-6-depleted embryos, this loss of asymmetry was associated with a pronounced increase in posterior PAR-3 density, and a smaller decrease in anterior PAR-3 density, relative to undepleted controls (Figure 7D; *cdc-42(RNAi)*, Movie S7). Thus, CDC-42 is required to maintain PAR-3 asymmetries in the absence of PAR-1, but it does not appear to do so simply through its ability to anchor and/or activate PAR-6/PKC-3 ^16,59^.

Depleting CDC-42 had no significant effect on PAR-3 oligomer disassembly rates (Figure 8A,B) or on the A/P ratio of fast release rates for monomeric PAR-3^V80D,D138K^::GFP (Figure S6M-P).

Asymmetric recruitment of PAR-3<u>^V80D,D138K^</u>::GFP was reduced, but not abolished, in embryos depleted of CDC-42 compared to controls (Figure 8F), even though endogenous PAR-3 remains asymmetrically enriched during maintenance phase in these embryos (Figure 7A,B). Reduced recruitment asymmetry cannot simply reflect reduced positive feedback on monomer recruitment, because this cannot explain the ***increase*** in posterior densities of PAR-3 observed in embryos depleted of CDC-42 (Figure 7D). An alternative possibility is that depletion of CDC-42 increases the basal (PAR-3 independent) recruitment rate, which would reduce the overall recruitment asymmetry if positive feedback on recruitment remains constant.

To test this possibility, we modified our model by reducing *k_basal_* to 50% of the control value while holding *k_f_* constant to approximate the recruitment asymmetry measured in CDC-42-depleted control embryos. Then we simulated PAR-3 dynamics from the initial distribution measured at late establishment phase in *par-1* homozygote embryos depleted of CDC-42 (Figure 7C,D; *cdc-42(RNAi)*). The simulations reproduced the qualitative signature and time course of PAR-3 asymmetry loss observed in CDC-42-depleted embryos, in which anterior PAR-3 levels fall and posterior levels rise toward an intermediate spatially uniform level (Figure 8G; 50% basal recruitment). Again, the loss of asymmetry was incomplete at late maintenance phase, as observed experimentally.

In summary, both PAR-6/PKC-3 and CDC-42 are required for dynamically stable PAR-3 asymmetries in the absence of posterior inhibition by PAR-1, but they suggest that PAR-6/PKC-3 and CDC-42 act in different ways: PAR-6/PKC-3 promotes positive feedback on PAR-3 recruitment, while CDC-42 reduces basal rates of recruitment, which indirectly increases the overall strength of positive feedback.

## Discussion

### Two feedback loops encode robust self-stabilizing unipolar PAR-3 asymmetries

Studies of PAR-mediated polarity have focused on how mutual antagonism between two sets of PAR proteins drive their stable enrichment in complementary cortical domains ^12,17,23,60–63^. However, the ability of PARs to form and stabilize *unipolar* asymmetries has remained poorly understood ^29–33^. Here we have combined single molecule analysis with mathematical modeling and experimental perturbations to identify two positive feedback loops that synergistically ensure robust, dynamically stable unipolar PAR-3 asymmetries in *C. elegans* zygotes. First, the intrinsic dynamics of PAR-3 membrane binding, oligomerization, and size-dependent oligomer dissociation encode negative feedback on PAR-3 dissociation. Second, PAR-3 acts indirectly to promote its own recruitment through a mechanism that depends on its conserved binding partners PAR-6/PKC-3. Using predictive mathematical models tightly constrained by empirically measured kinetics, we have shown that these two modes of positive feedback are individually necessary and jointly sufficient to account, both qualitatively and quantitatively, for the locally inducible and dynamically stable PAR-3 asymmetries observed in zygotes lacking posterior inhibition by PAR-1. Thus, a tight correspondence between model-predicted and observed dynamics, integrating multiple independent measurements of single molecule kinetics, oligomer size distributions and the time evolution of cell-scale asymmetries, reveals a robust, internally self-consistent and quantitively accurate picture of how self-stabilizing unipolar PAR-3 asymmetries emerge from molecular scale kinetics.

PAR-3 is the keystone member of the PAR polarity network, which is required for all other PAR asymmetries. In the *C. elegan*s zygote, PAR-3 is the only member that remains asymmetric when all other known members of the PAR network are absent or symmetrically distributed ^17^. In the *C. elegans* zygote, and in other cells, the spatial distribution of PAR-3 determines where PAR-6/PKC-3 load onto the membrane and into a complex with active CDC-42, to form a gradient of PKC-3 activity that defines anterior (or apical) identity and opposes posterior (or basolateral) identity ^16,17,64^. Thus, the spatial distribution of PAR-3 dictates polarity, even in cells where mutual inhibition plays an important role.

Our analysis of a model tightly constrained by empirical measurements reveals that the combination of negative feedback on PAR-3 dissociation and positive feedback on PAR-3 recruitment encodes the co-existence of two dynamically stable states: a spatially uniform state with high PAR-3 levels, as observed during maintenance phase in zygotes that fail to polarize due to lack of a functional sperm cue; and a polarized state with high anterior and low posterior levels of PAR-3, matching those observed during maintenance phase in polarized zygotes.

Importantly, the model predicts that local depletion of PAR-3, mimicking the effects of cortical flow or local PAR-1 accumulation during zygotic symmetry breaking ^11,12^, can induce an irreversible transition from the spatially uniform to the stably polarized state. Thus, the intrinsic kinetics of PAR-3 membrane-binding and oligomerization, plus weak feedback on PAR-3 recruitment mediated by PAR-6/PKC-3, endow *C. elegans* zygotes with a robust symmetry-breaking response to the sperm cue, independent of mutual antagonism. Because these core features of PAR-3 and its interactions with PAR-6/PKC-3 are conserved across the metazoa, it seems likely that they contribute to stabilizing unipolar asymmetries observed in other contexts, for example in neuroblasts ^27,31^ or meiotic oocytes^65^, and that they may play key roles in controlling PAR asymmetries, even in cells like the *C. elegans* zygote that also rely on mutual inhibition.

### Intrinsic dynamics of membrane binding and oligomerization encode positive feedback on PAR-3 accumulation

Self-oligomerization is a conserved property of PAR-3^34^. Recent studies in *C. elegans* zygotes have emphasized how this property can contribute to the advective transport of PAR-3, and its binding partners PAR-6/PKC-3, during symmetry-breaking, through increased physical coupling to the cortical actomyosin cytoskeleton and through increased residence time via size-dependent binding avidity ^17,26,39–41^. Here we have shown that the intrinsic dynamics of membrane binding and oligomerization also confer positive feedback on PAR-3 accumulation. Our model predicts, and our experiments confirm, a positive dependence of mean oligomer size on PAR-3 density, and we have measured directly a sharp decrease in oligomer dissociation rate with size.

Together these make the average rate constant for PAR-3 dissociation a sharply decreasing function of PAR-3 density. For measured rates of size-dependent oligomer dissociation (Figure 3K), oligomer disassembly (*k_diss_* = ∼ 0.1 *sec*), and monomer dissociation (*k_off_* = ∼3 *sec*), our model predicts that this negative feedback on PAR-3 dissociation contributes a nearly 4-fold A:P difference in the flux of PAR-3 molecules off the membrane, accounting for half the total asymmetry observed in *par-1* mutant embryos lacking posterior inhibition of PAR-3.

Negative feedback on dissociation is a generic consequence of reversible membrane binding and oligomerization. In this simple generic form, it does not consume energy, and therefore it cannot alone sustain dynamically stable asymmetries. However, it can sharply reduce the amount of additional energy-consuming feedback needed for stable asymmetries. Because weak membrane binding and self-oligomerization are widespread and easily evolved properties of proteins, we expect that many types of cells use this form of feedback to promote symmetry-breaking and stable polarity. Indeed, our results add to a growing body of work emphasizing the importance of clustering/oligomerization of peripheral membrane binding proteins to the establishment and maintenance of cell surface asymmetries in polarized cells^17,66–68^.

### PAR-6/PKC-3 mediate positive feedback on PAR-3 monomer recruitment

Our observations suggest that positive feedback on PAR-3 monomer recruitment supplies the additional feedback required for dynamically stable PAR-3 asymmetries. First, oligomerization-defective PAR-3<u>^V80D,D138K^</u>::GFP binds at > 2-fold higher rates to anterior membranes where endogenous PAR-3 itself is enriched. In principle, this binding asymmetry could reflect either a fixed PAR-3-independent bias, or positive feedback on monomer recruitment. However, a mathematically controlled comparison reveals that positive feedback on monomer recruitment, and not a fixed recruitment bias, can explain not only the observed dependence of PAR-3 asymmetry on total PAR-3 density, but also the complete loss of PAR-3 asymmetry induced by a partial reduction of total PAR-3 (Figure 6D).

Our experiments identify a central role for PAR-6/PKC-3 in stabilizing PAR-3 asymmetries through positive feedback on PAR-3 monomer recruitment. Embryos co-depleted of PAR-6 and MRCK-1 establish PAR-3 asymmetries initially, but these are rapidly lost during maintenance phase. Co-depleting embryos of PAR-6 and MRCK-1 completely blocks the asymmetric recruitment of PAR-3<u>^V80D,D138K^</u>::GFP in early maintenance phase, even though endogenous PAR-3 remains highly asymmetric at this stage, but such co-depletion has minimal effects on PAR-3 oligomer disassembly or monomer dissociation rates. Finally, a variant of our empirically-constrained model lacking positive feedback on monomer recruitment correctly predicts the mode and time course of asymmetry loss observed in experiments. Importantly, acute inhibition of PKC-3 kinase activity also blocks asymmetric recruitment of PAR-3<u>^V80D,D138K^</u>::GFP in early maintenance phase. Thus, PAR-6/PKC-3, and PKC-3 kinase activity are required for positive feedback on PAR-3 monomer recruitment, and this feedback is essential for dynamically stable unipolar PAR-3 asymmetries in the absence of posterior inhibition by PAR-1.

Strikingly, although PAR-6/PKC-3 fulfill many of their roles in polarity as part of an active complex with CDC-42, they appear to function independently of CDC-42 to promote positive feedback on PAR-3 monomer recruitment. Whereas loss of PAR-6/PKC-3 reduces both anterior and posterior accumulation of PAR-3 and abolishes asymmetric recruitment of PAR-3<u>^V80D,D138K^</u>::GFP, CDC-42 depletion ***increases*** both anterior and posterior accumulation of PAR-3, and thereby ***reduces*** asymmetric recruitment of PAR-3<u>^V80D,D138K^</u>::GFP. Our modeling confirms that both of these effects of depleting CDC-42 can be explained by an increase in the basal rate of PAR-3 monomer recruitment, suggesting the that CDC-42 may have an independent role in controlling the availability of cytoplasmic PAR-3 monomers for membrane binding.

How PAR-6/PKC-3 might act independently of CDC-42 to mediate positive feedback on PAR-3 recruitment remains unclear, but previous studies and our data constrain the class of possible mechanisms. PAR-3 asymmetries persist during maintenance phase in embryos co-depleted of CHIN-1 and PAR-1 in which active CDC-42, PAR-6 and PKC-3 are uniformly enriched ^17^.

Thus, positive feedback on PAR-3 monomer recruitment does not require the asymmetric enrichment of PAR-6/PKC-3. Instead, we propose that local interaction of PAR-6/PKC-3 with PAR-3 induces a change in PAR-6/PKC-3 activity state which licenses their ability to recruit additional PAR-3 monomers from the cytoplasm. A similar mechanism has been proposed for positive feedback on Rho GTPase activation by Anillin ^69^.

One possibility is that PAR-6/PKC-3 dimers bound to PAR-3 could recruit a second PAR-3 molecule. Strong binding of PAR-6/PKC-3 to PAR-3 *in vitro*, and membrane recruitment of PAR-6/PKC-3 *in vivo*, requires direct binding between a PDZ-binding motif (PBM) on PKC-3 and the PDZ2 domain of PAR-3 ^54^. However, PAR-6/PKC-3 heterodimers can also bind PAR-3 via PAR-6 ^52^, or through an interaction between the kinase domain of PKC-3 and PAR-3 ^54,70^.

Thus, in principle, PAR-6/PKC-3 heterodimers complexed with PAR-3 at the membrane could recruit additional molecules of cytoplasmic PAR-3. Indeed, a recent study suggests that multivalent binding of PAR-6/PKC-3 to PAR-3 mediates the cooperative assembly of PAR-3 oligomers during polarity establishment phase ^71^. For this to constitute positive feedback, an energy-consuming step (e.g., phosphorylation of PAR-3 by PKC-3) would be required. Moreover, the fast mobilities of newly-bound PAR-3<u>^V80D,D138K^</u>::GFP molecules imply that their asymmetric recruitment does not require direct stable binding to existing PAR-3 oligomers. Thus, a bridging mode of recruitment would have to be very transient, delivering cytoplasmic monomers into a freely diffusing, membrane-bound state, or it would have to prefer the use of membrane-bound PAR-3 monomers over oligomers. Consistent with this latter possibility, our observation that PAR-3<u>^V80D,D138K^</u>::GFP remain asymmetrically recruited in embryos lacking endogenous PAR-3 (Figure 8E), suggests that membrane-bound monomeric PAR-3 is sufficient for positive feedback on monomer recruitment.

An alternative possibility is that PAR-3 could promote direct binding of PAR-6/PKC-3 heterodimers to the plasma membrane via PKC-3’s C1 domain. The C1-membrane is autoinhibited through an intramolecular association with the catalytic domain, and release of this inhibition is thought to require binding to PAR-3 and PAR-6 ^54,72^. Thus, in principle, PAR-3 could act to locally recruit PAR-6/PKC-3 heterodimers into a membrane-bound state, which could in turn recruit additional PAR-3 molecules. Again, for this to constitute feedback, an energy-consuming step would be required. Distinguishing these and other possible mechanisms for PAR-6/PKC-3-mediated feedback on PAR-3 monomer recruitment will be an important goal for future work.

## Supporting information

Modeling Supplement

## Acknowledgments

This work was funded by NIGMS grants T32 GM007197 and 1R01GM098441.

---

**Supplementary Figure S1 (related to Figure 2).**
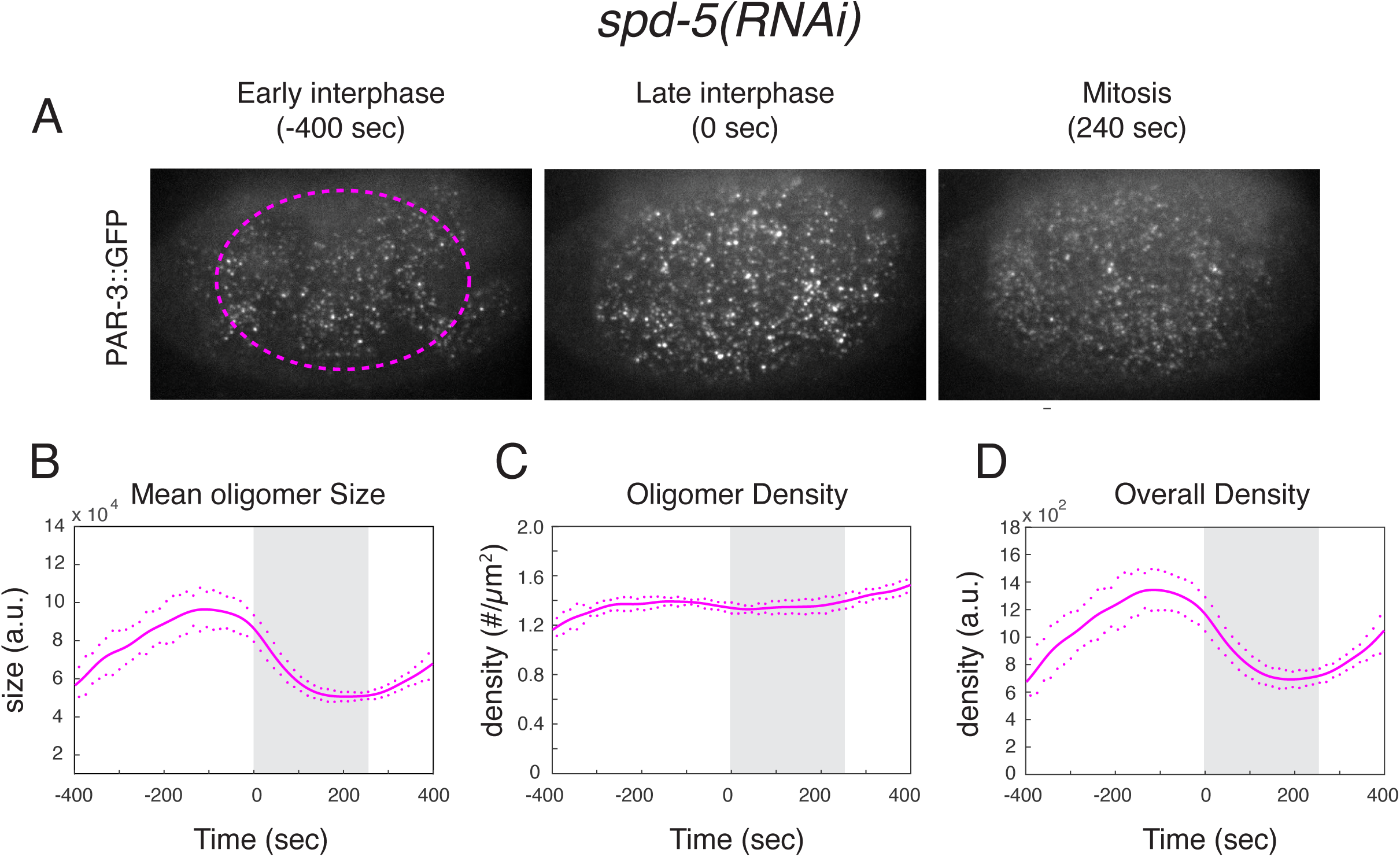
PAR-3 oligomer dynamics during maintenance phase in *spd-5(RNAi)* embryos. **(A)** Individual near-TIRF images from a time-lapse sequence showing PAR-3 oligomer intensities and distributions at the cell surface in *spd-5(RNAi)* embryos at early establishment, late establishment, and late maintenance phase. **(B-D)** Results of particle detection analysis showing **(B)** mean oligomer size, **(C)** oligomer density, and **(D)** total PAR-3 fluorescence density over time on the cortex. Data are from n = 6 embryos aligned with respect to the onset of maintenance phase. The grey shaded area indicates maintenance phase. Solid lines indicate the mean and dots indicate SEM at each time point.

**Supplementary Figure S2 (related to Figure 3 and 5).**
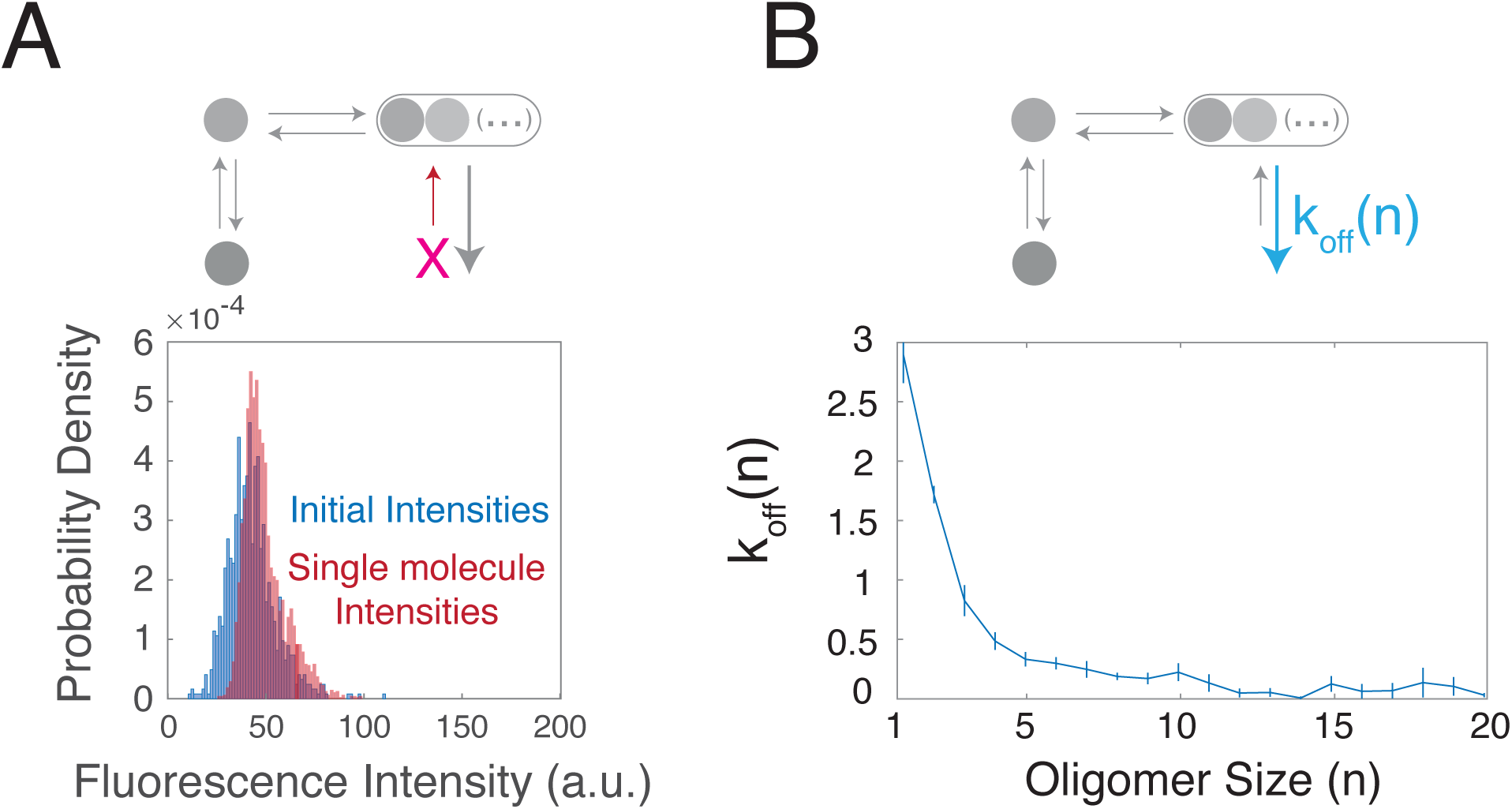
Single molecule measurements of size-dependent recruitment and dissociation of PAR-3::GFP oligomers. **(A)** Overlay of the distributions of background-subtracted intensities of (blue) newly detected PAR-3::GFP speckles from the data in (A) and (peach) single molecule speckles after photobleaching in the same cells. **(B)** Estimated rate constants for dissociation rates of PAR-3::GFP oligomers from the membrane as a function of inferred number of subunits n, measured as described for mNG::PAR-3 in Figure 3K. Error bars represent the SEM. Data were compiled from n = 6 individual embryos.

**Supplementary Figure S3 (related to Figure 4).**
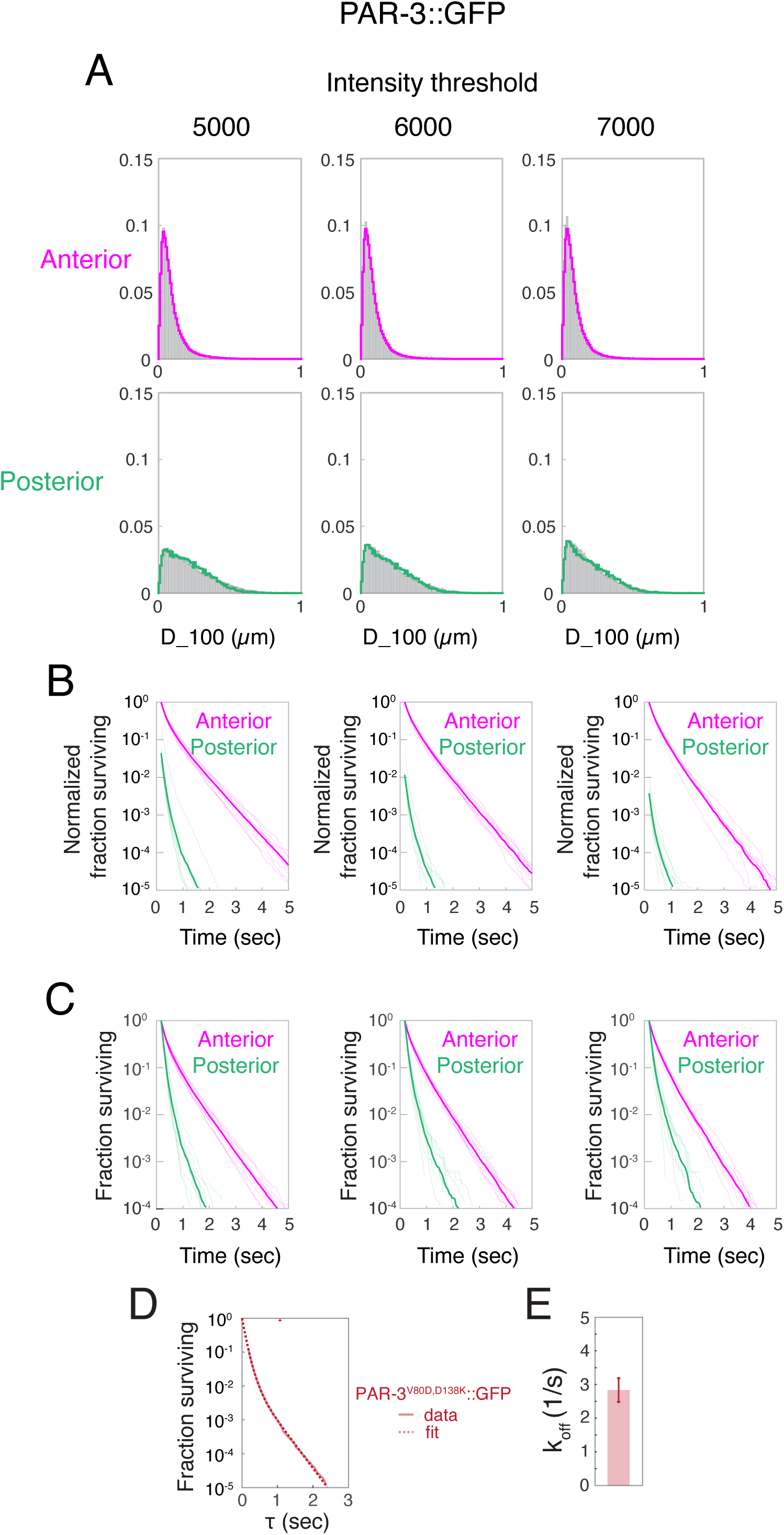
Measured PAR-3 release rates and mobilities do not depend sensitively on single molecule detection threshold. **(A)** Distributions of single molecule displacements after 100 msec (D_100) measured for endogenously-tagged PAR-3::GFP molecules tracked on the anterior and posterior cortex in n = 9 embryos. Each column represents data obtained using a different integrated intensity threshold for detecting single molecules. Solid curves show least squares best fit to weighted D_100 distributions measured for oligomers and for simulated Brownian diffusion with D = 0.1 µm^2^/sec, as in Figure 4.. **(B)** Release curves measured for single molecules of PAR-3::GFP on the anterior and posterior cortex of wild type embryos under continuous illumination for different intensity thresholds. Thin traces show release curves for individual embryos (n = 9); Thick traces show pooled data for all embryos. Data were normalized by total numbers of anterior tracks. **(C)** Same data plotted in **(B)**, but with anterior and posterior data individually normalized to emphasize differences in release kinetics. **(D)** Release curves measured for posterior single molecules of oligomerization-defective PAR-3. Dashed lines indicate data fit to a weighted sum of two exponentials. **(E)** Corresponding estimate of effective *k_off_* from the data in (D). n = 11 embryos. Error bars represent 95% confidence intervals obtained by bootstrap sampling.

**Supplementary Figure S4 (related to Figure 5).**
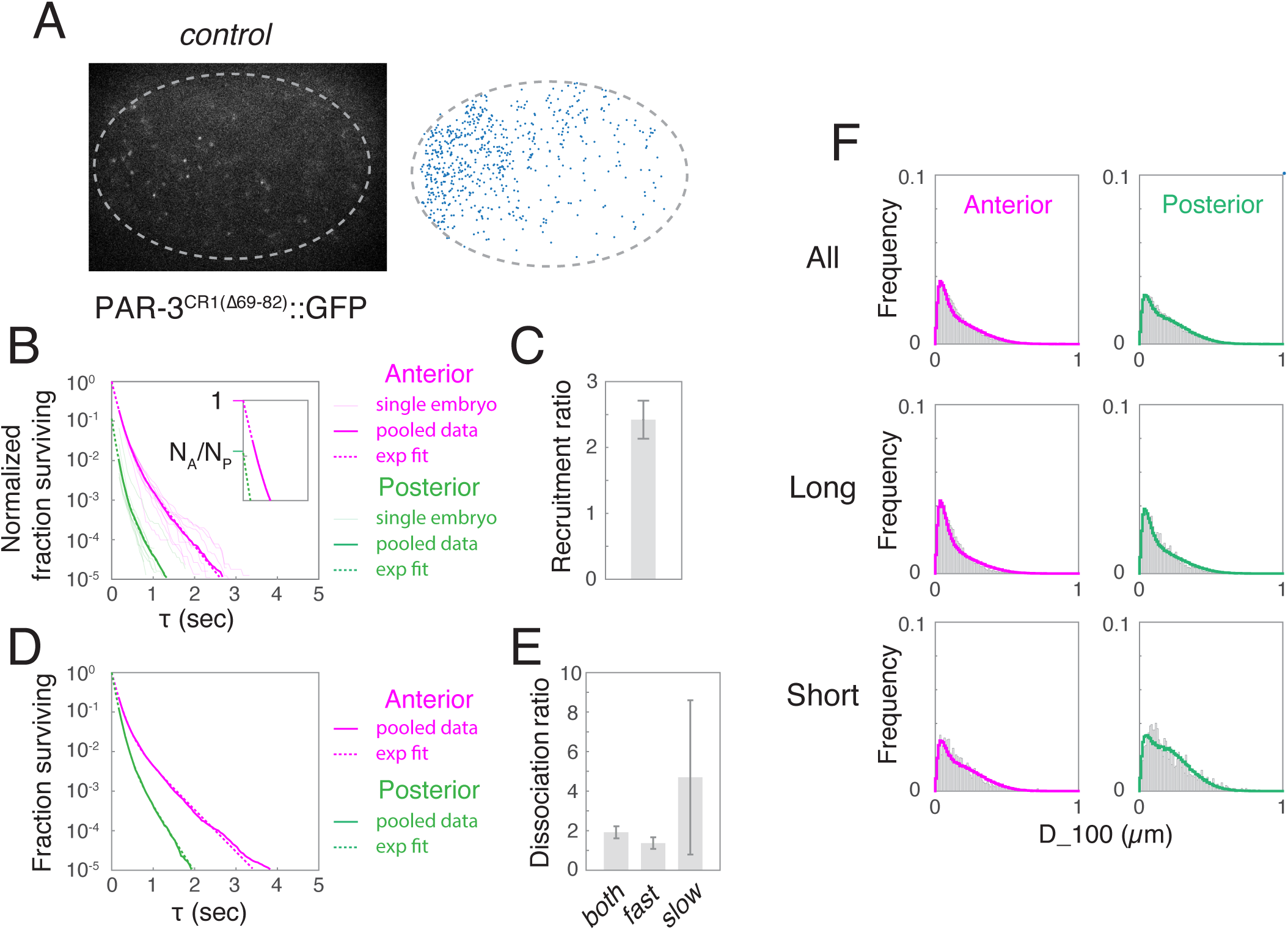
Single molecule analysis of monomer recruitment and release rate asymmetries for a second form of oligomerization-defective PAR-3 (PAR-3^CR1Δ(69–82)^::GFP) (A) Representative images and distribution of single molecule appearance events for oligomerization-defective PAR-3^CR1Δ(69-82)^::GFP, detected by single molecule imaging. Each blue dot represents a single binding event. **(B)** Release curves for (n = 10) embryos. Faint lines show data for individual embryos, thick lines for pooled data, and dashed lines indicate the exponential fits to pooled data. Inset shows the extrapolated zero-crossings from which the total numbers of binding events were inferred. Anterior and posterior data were normalized to total numbers of anterior binding events to highlight the recruitment asymmetry. **(C)** A:P recruitment ratios measured from the data in **(B)**. Error bars indicate 95% confidence intervals obtained from bootstrap sampling. **(D)** Pooled release curves and exponential fits with anterior and posterior data separately normalized to highlight differences in release rates. **(E)** A:P ratios of monomer release rates for fast and slow components (both) and for fast components alone (fast). Error bars indicate 95% confidence intervals obtained from bootstrap sampling. **(F)** Distributions of short-term displacements (D_100) for anterior and posterior molecules, measured over all trajectories (top row), long trajectories (middle row; duration > 2 sec), and short trajectories (bottom row; duration = 250 msec). Solid curves show data fits to weighted sums of D_100 distributions for oligomers and for simulated Brownian diffusion with diffusivity D = 0.1 µm^2^/sec (as in Fig 4C).

**Supplementary Figure S5 (related to Figure 6).**
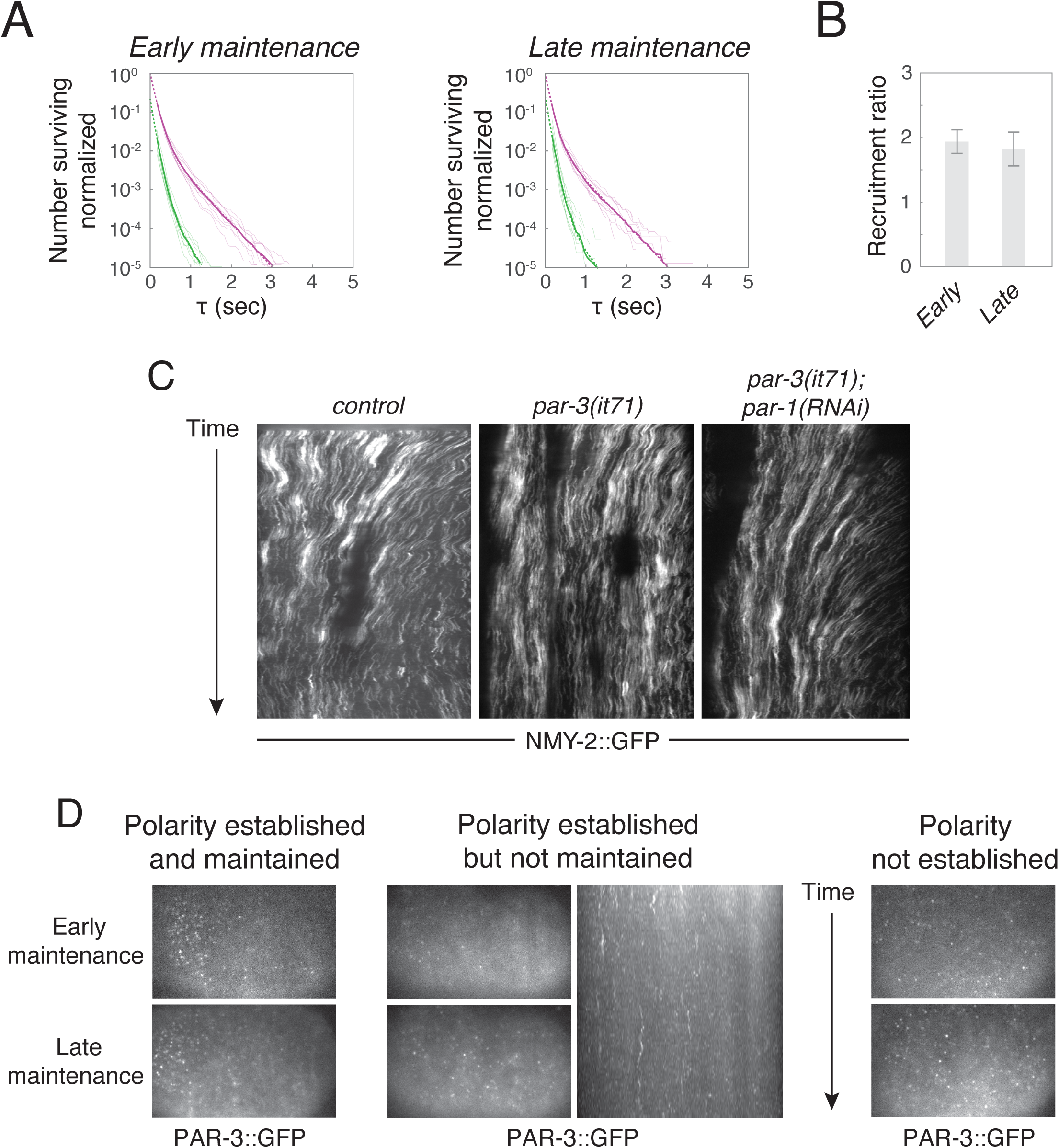
Further details underlying the model and its comparison to data. **(A)** (left) Release curves for PAR-3^V80D,D138K^::GFP molecules measured in the same embryos during early and late maintenance phase. Anterior and posterior data were normalized to total numbers of anterior binding events to highlight the recruitment asymmetry. (right) A:P recruitment ratios measured from the same data (n = 10 embryos). Error bars indicate 95% confidence intervals obtained from bootstrap sampling. **(B)** Kymographs illustrating patterns of cortical flow during establishment phase in control embryos, *par-3* mutant embryos, and *par-3* mutant embryos depleted of PAR-1 by RNAi. **(C)** Examples of different phenotype classes in the PAR-3 depletion experiment shown in Figure 5D. Top and bottom micrographs show the distribution of PAR-3 at early and late maintenance respectively. We observed three distinct classes: (left) polarity established and maintained, (middle) polarity established and not maintained and (right) polarity never established. For the middle class, the kymograph confirms that the loss of PAR-3 asymmetry was not due to redistribution of PAR-3 by posterior-directed cortical flows.

**Supplementary Figure S6 (related to Figure 8).**
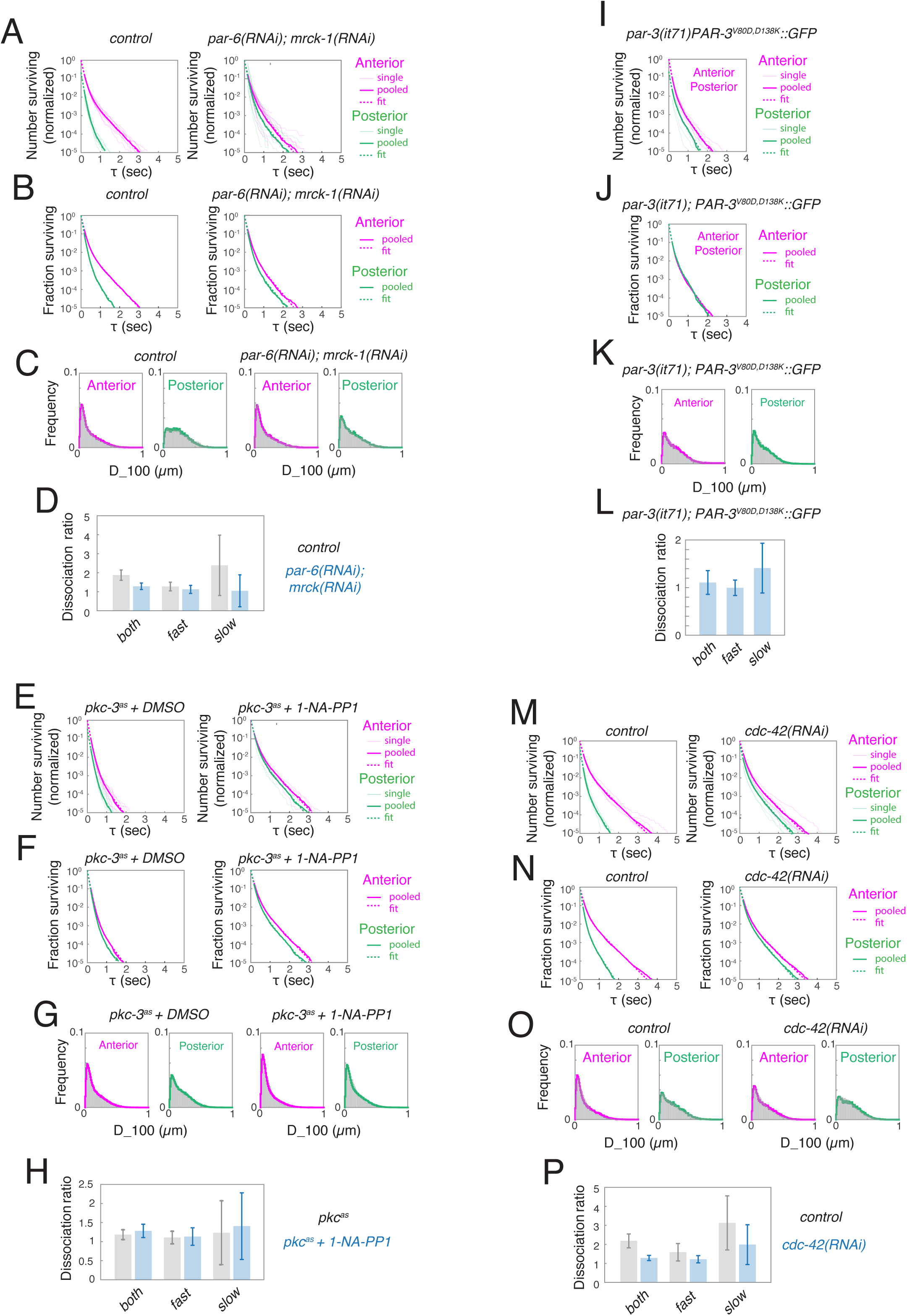
Mobilities and release curves for single molecules of oligomerization-defective PAR-3^V80D,D138K^::GFP under different perturbations. (A,E,I,M) Release curves for PAR-3^V80D,D138K^::GFP molecules in: (A) control (n = 9) and *par-6(RNAi);mrck-1(RNAi)* (n = 12) embryos, (E) *pkc-3^as^* embryos treated with DMSO (n = 6) or 50 µM 1-NA-PP1 (n = 8), (I) *par-3* mutant embryos partially rescued by transgenic PAR-3^V80D,D138K^::GFP (n = 6), and (M) control (n = 6) and *cdc-42 (RNAi)* (n = 6) embryos. Faint lines show data for individual embryos, thick lines for pooled data, and dashed lines indicate the exponential fits to pooled data. Anterior and posterior data were normalized by total numbers of anterior binding events to highlight the recruitment asymmetry. (B,F,J,N). The same data with anterior and posterior release curves normalized using total numbers of anterior and posterior binding events respectively. (C,G,K,O). Distributions of short-term displacements (D_100) for the indicated genotypes and perturbations. Solid curves show data fits to weighted sums of D_100 distributions for oligomers and for simulated Brownian diffusion with diffusivity D = 0.1 µm<u>^2^</u>/sec (as in Fig 4C). Recruitment ratios computed from these data are shown in Figure 8. **(D,H,L,P)**. Dissociation ratios measured for the indicated genotypes and perturbations. **(both):** ratio of effective release rates for slow and fast dissociating monomers. **(fast):** Estimated ratio of release rates for fast-dissociating monomer fraction. **(slow):** Estimated ratio of release rates for slow-dissociating monomer fraction. Error bars represent 95% confidence intervals obtained by bootstrap sampling.

## Supplemental Movie Legends

**Movie S1. Dynamics of endogenously-tagged PAR-3 during polarity establishment and maintenance phases.** Left: embryo from *par-1* heterozygote mother; Middle: Embryo from *par-1* homozygote mother; Right: *spd-5(RNAi)* embryo. All three embryos were imaged under identical conditions. Time is measured relative to the onset of maintenance phase. Time compression 90:1

**Movie S2. Dynamics of FRAP recovery**. Embryo was subjected to photobleaching just before time 0. Time compression 18:1.

**Movie S3.** Fast imaging of endogenously-tagged mNeonGreen::PAR-3 in an embryo depleted of myosin II heavy chain (*nmy-2(RNAi)*). Time compression 1:1.

**Movie S4. Single molecule imaging of endogenously-tagged PAR-3::GFP.** Data were collected with high laser power in streaming mode at 20 frames per second to measure short-term mobilities and monomer dissociation rates. Time compression 1.5:1.

**Movie S5.** Single molecule observations of appearance events in control and *par-1(RNAi)* embryos expressing transgenic PAR-3::GFP. (left) *control*; (right) *par-1(RNAi)*. Data were collected with high laser power in streaming mode at 20 frames per second to measure monomer recruitment and dissociation kinetics and short-term mobilities. Time compression 1.5:1.

**Movie S6.** Distribution of endogenously-tagged PAR-3::GFP oligomers during maintenance phase in embryos from *par-1*/+ mothers subjected to the indicated RNAi depletions. Time is measured relative to the onset of maintenance phase. Time compression: 90:1

**Movie S7.** Distribution of endogenously-tagged PAR-3::GFP oligomers during maintenance phase in embryos from *par-1*/*par-1* mothers subjected to the indicated RNAi depletions. Time is measured relative to the onset of maintenance phase. Time compression: 90:1

## STAR Methods

### C. elegans culture and RNAi

We cultured *C. elegans* strains under standard conditions ^73^ on 60mm petri plates containing *E. coli* bacteria (OP50) raised on normal growth medium (NGM). Table 1 provides a list of strains used in this study. Unless otherwise specified, strains were provided by the Caenorhabditis Genetics Center, which is funded by the National Center for Research Resources.

**Table 1.** *C. elegans* strains used in this study.

| Name | Genotype | Source |
| --- | --- | --- |
| KK1216 | par-3(it298[PAR-3::GFP]) III | CGC |
| KK292 | rol-4(sc8) par-1(it0051) V/DnT1 IV;V | Ken Kemphues |
| EM292 | par-3(it298[PAR-3::GFP]) III; rol-4(sc8) par-1(it0051) V/DnT1 IV;V | This study; made from KK1216 and KK292 |
| KK1039 | itIs268[ppar-3::PAR-3(V80D D198K)::GFP + unc-119(+)]; unc-119(ed3) III | Ken Kemphues |
| KK973 | itIs169 [ppar-3::PAR-3 CR1 delta(69-82), unc-119(+)]; unc-119(ed4) III | Ken Kemphues |
| KK1038 | itIs268[ppar-3::PAR-3(V80D D198K)::GFP + unc-119(+)]; par-3(it71) unc-32(e189) III | Ken Kemphues |
| LP232 | par-6(cp60[par-6::mKate2::3xMyc + LoxP unc-119(+) LoxP]) I par-3(cp54[mNG::3xFlag::par-3]) III; | Daniel Dickinson |
| NWG316 | <u>*pkc-3(crk77[l331A,T394A]) II; par-2(it328[gfp::par-2]) III*</u> | CGC |
| EM547 | *pkc-3(crk77[l331A,T394A]) II*; itIs268[ppar-3::PAR-3(V80D D198K)::GFP + unc-119(+)]; unc-119(ed3) III | This study; made from NWG316 and KK1039 |
| KK1060 | itIs192[pie::GFP::PAR-3 unc119(+)]; (itIS272[pie-1::mCherry::PAR-6 unc119(+)] | Ken Kemphues |
| EM17 | lon-1(e185) PAR-3(it71)/qC1 dpy-19(e1259) glp-1(q339)III; zuIs45 [pnmy-2::NMY-2::GFP + unc-119(+)] V | This study |

We performed RNAi experiments using the feeding method ^74^. We obtained bacteria targeting *par-1*, *par-3*, *par-6*, *cdc-42*, *mrck-1*, *spd-5*, *perm-1* and *nmy-2* from the Kamath RNAi library ^75^, and bacteria targeting GFP from Jeremy Nance. Briefly, we grew bacteria to log phase in LB with 50µg/ml ampicillin, seeded ∼300µl of culture onto NGM plates supplemented with 50µg/ml ampicillin and 1mM IPTG, incubated plates at room temperature for two days, and then stored them at 4°C for up to one week before use. We transferred L4 larvae or young adults (for *par-3(RNAi*)) to feeding plates and then cultured them at 25°C before imaging as follows: 0 - 8 hours at 25°C for *par-3(RNAi*) and 24 – 48 hours at room temperature (21-23°C) for all others, adjusting exposure time to achieve the desired phenotype. We verified strong depletions of SPD-5 by observing complete failure of polarity establishment, strong depletions of PAR-6 and CDC-42 by observing symmetric first and second cleavages, and strong depletions of NMY-2 and MRCK-1 by observing absence of cortical flows during polarity establishment and/or maintenance phases.

### Inhibition of analog-sensitive PKC-3 activity

We inhibited PKC-3 activity as previously described ^76^. Briefly, we placed worms expressing oligomerization-defective PAR-3 (PAR-3(V80D D198K)::GFP) and the analog-sensitive PKC-3 on perm-1(RNAi) plates for 16-20 hours to permeabilize the eggshells. We cut open worms into embryonic culture medium (ECM: 10% 5mg/ml Inulin, 20% 250mM HEPES pH 7.4, 20% heat-inactivated FBS, 50% Leibowitz L-15 medium) containing 50 µM ATP analog 1-NA-PP1 or DMSO vehicle control. We transferred the embryos to 2% agarose pads prepared using ECM (without FBS) and either 1-NA-PP1 or DMSO vehicle control, and then performed imaging and analysis as described below. We verified successful inhibition of PKC-3 by observing symmetric first cleavage.

### Live imaging

For all other experiments, we mounted embryos in standard eggs salts on 2% agarose pads. For data reported in Figures 3-8 and S2-S6, we used an Olympus IX50 inverted microscope equipped with an Olympus OMAC two-color TIRF illumination system, a CRISP autofocus module (Applied Scientific Instrumentation), and a 1.49 NA oil immersion TIRF objective. Laser illumination at 488 nm from a 50-mW solid-state Sapphire laser (Coherent) was delivered by fiber optics to the TIRF illuminator. We magnified images by 1.6x and collected them on an Andor iXon3 897 EMCCD camera, yielding a pixel size of 100 nm. Image acquisition was controlled by Andor IQ software. For the data reported in Figures 2, 7 and S1, we used a Nikon Ti2-E inverted microscope equipped with a LunF XL laser combiner housing solid state 488nm, 561nm and 640nm lasers and feeding three separate TIRF illumination arms, a Ti2 N-ND-P Perfect Focus unit, and a CFI APO 100X 1.49NA TIRF lens. We magnified images by 1.5X and collected them on an Andor IXON Life 897 EM-CCD camera to achieve a pixel size of ∼106nm. Image acquisition was controlled by Nikon Elements software.

For all experiments, we used near-TIRF illumination. We set the laser illumination angle to standard values that we chose empirically to approximately maximize signal-to-noise ratio while maintaining approximately even illumination across the field of view. All embryos were oriented with their AP axis perpendicular to the axis of TIRF illumination to avoid any confounding effects of illumination gradients in comparisons of anterior vs posterior PAR-3 dynamics. Further details of the imaging conditions used for specific quantitative analyses appear below.

### Image analysis

We performed particle detection and tracking as previously described ^15,17^. Briefly, we used a slightly modified Matlab implementation (http://people.umass.edu/kilfoil/downloads.html) of methods introduced by Crocker and Grier ^77^ to detect diffraction-limited features of interest as local intensity peaks in bandpass filtered images and then to determine their centroids and background-subtracted integrated intensities. We used µTrack software (http://github.com/DanuserLab/u-track) developed by Jaqaman and colleagues ^78^ to track features over time. We integrated particle detection and tracking and all subsequent analyses using custom MATLAB scripts that are available upon request.

### Timelapse analysis of PAR-3 oligomer dynamics

We measured PAR-3 oligomer dynamics in *par-1(it51)* mutant embryos expressing a C-terminal fusion of GFP to PAR-3 from the endogenous locus (hereon *par-1*; PAR-3::GFP), using *par-1/+* heterozygotes as controls. We collected images using near-TIRF illumination in timelapse mode using 10% laser power and 50 msec exposure times at 3 sec intervals to minimize photobleaching. For each embryo and each timepoint, we identified PAR-3 clusters within hand-drawn anterior and posterior regions of interest (ROIs). For this and all subsequent measurements of feature intensities, we measured background-subtracted intensities by measuring integrated intensity within a circular ROI with radius of 3 pixels (300 nm) around the feature’s centroid position and using the mean intensity within an annulus of width 2 pixels surrounding the central ROI to estimate local background. We then determined the minimal polygon containing all detected features and used its area to compute the densities of features or total PAR-3 fluorescence. For *spd-5(RNAi)* embryos, we used a single ROI for all measurements. For individual embryos, we plotted the smoothed mean intensities vs time to determine the timepoint in late establishment phase when mean intensities began to fall sharply, which coincided roughly with the onset of pseudocleavage (PC) furrow relaxation. Then we used this timepoint to align data across multiple embryos. We then computed means and standard errors across embryos from un-normalized data.

### FRAP experiments

We performed FRAP experiments using the following protocol: We collected 10 images with 50 msec exposures using near-TIRF illumination at 1 second intervals using 30% laser power. We then subjected embryos to constant near-TIRF illumination with 100% laser power for 1.5 seconds to photobleach molecules at the cell surface. We then tracked recovery in timelapse mode with 50 msec exposures at 3 second intervals using 30% laser power. Because PAR-3 levels decrease during early maintenance phase, we normalized measurements of fluorescence during recovery using unbleached control data collected over the same timeframe.

### Measurements of PAR-3 oligomer size distribution

We measured the distribution of PAR-3 oligomer sizes during late maintenance phase in homozygous *par-1*; PAR-3::GFP mutant embryos. We first identified embryos in establishment phase before pseudocleavage. We used 30% laser power and 50 msec exposures at 1 sec intervals to focus on the cortex while minimizing photobleaching. Following pseudocleavage relaxation, we waited for 4 minutes and then acquired images in streaming mode for 50 seconds using 100% laser power and a 50 msec exposure time. For each embryo, we used hand-drawn ROIs to separate anterior and posterior features. We pooled all features detected in the first three frames and used these to estimate the distribution of PAR-3::GFP oligomer intensities. During the last 20 seconds of continuous exposure to 100% laser power, the distribution of background-subtracted feature intensities approximated a Gaussian distribution, as described previously ^15^. We took the mean of a gaussian fit to this distribution as our estimate of single molecule intensities.

We compared three different approaches to estimating the distribution of PAR-3 oligomer sizes. In the first approach, based on Bayesian inference, we initially assumed each feature had an equal prior probability of containing between 1 and 20 subunits. We then calculated the posterior probabilities that an observed feature contained n subunits (0 < n <=20) given the conditional probabilities of an n-mer producing the observed fluorescence intensity, assuming that means and variances for the distributions of n-mer intensities were equal to the mean and variance of the single molecule distribution multiplied by n. Then we used these posterior probabilities to update the prior probabilities, repeating until the prior and posterior probability distributions converged, yielding an estimate of the steady-state size distribution. In a second approach, based on maximum likelihood estimation, we assumed that the distribution of feature intensities equals a weighted sum of the distributions of n-mers of size 1 to 20, determining the distributions of n-mer intensities as described above. We then used the lsqnonlin function in MATLAB to compute the most likely values for each of the weights. Finally, as a third approach, we binned feature by intensity, assigning all feature with intensities in the interval [(n – 0.5)*I_sm, (n+0.5)*I_sm)] to size bin n, where I_sm is the mean intensity of single GFP molecules. Because all three approaches produced very similar results, we used the third (simplest) approach to produce the size distributions shown in Figure 3.

### Measuring oligomer disassembly

To measure oligomer disassembly rates K_dis_, we used a transgenic strain expressing PAR-3::GFP. We used RNAi against GFP followed by photobleaching to reduce the density of transgenic PAR-3::GFP to single molecule levels as described previously ^15^. We then imaged single PAR-3 molecules using 100 % laser power and 50 msec exposures, holding photobleaching rate *k_pb_* constant, while varying the duty ratio (dr) of exposure, such that, for each duty ratio, the effective photobleaching rate is given by *k_pb_dr*. For each duty ratio, we performed single particle tracking and constructed release curves as described above, pooling the trajectories from all embryos. We fit the release curves to exponential functions of the form x(t) = x_0_e^-Kobst^, using tmin = 1 sec to exclude the more rapid (∼3/sec) monomer dissociation.

We then used MATLAB’s built-in ***fit*** function to obtain a linear least squares fit to K_obs_ = K_dis_ + *dr* ∗ *k_pb_*. We took the y-intercept and slope of the linear fits as our estimates of K_dis_ and *k_pb_*, respectively, and computed confidence intervals for the estimate using bootstrap sampling of the pooled trajectories To compare oligomer disassembly rates in control, *par-6(RNAi);mrck-1(RNAi)* and *cdc-42(RNAi)* embryos, we imaged all embryos for 120 seconds during early maintenance phase with a 5% duty ratio, and the same fixed unit exposure as above (50 msec at 100% laser power). We constructed release curves and measured K_obs_ for each embryo as described above using tmin = 2 sec. Then we used the estimate of *k_pb_* from above to estimate K_dis_ = K_obs_ − *dr* ∗ *k_pb_* for each embryo.

### Measuring oligomer size-dependent dissociation

To measure oligomer size-dependent dissociation rates, we imaged embryos expressing NeonGreen::PAR-3 or PAR-3::GFP in streaming acquisition mode using 30% laser power and 50 *msc* exposure times during maintenance phase in embryos depleted of NMY-2. We performed particle tracking on the first 20 frames of each image sequence to avoid confounding effects of photobleaching on oligomer size inference. We inferred the size of each tracked particle based on its initial intensity using the single molecule distribution as a standard, as described above. For each inferred oligomer size n, we computed the fraction of oligomers F_*n*_ that remained after 1 sec. Assuming a single-rate first-order dissociation reaction, we solved F_*n*_ = e^-Koff(*n*)^ to estimate *k_off_*(*n*) = −ln (F_*n*_).

### Measuring single molecule recruitment ratios, dissociation rates and mobilities

We measured single molecule recruitment ratios, unbinding rates and mobilities from data collected in near-TIRF mode using streaming acquisition with the same “unit exposure” (50 msec exposure times and 100% laser power) used to measure oligomer disassembly and photobleaching rates above. We used a brief (∼5 sec) pre-exposure to 100% laser power in widefield mode to reduce the density of labeled molecules, and then allowed embryos to equilibrate surface densities of labeled molecules for > 60 secs before acquisition. We then collected single molecule data and performed particle tracking analysis as described above. For each embryo, we used hand-drawn ROIs to isolate anterior and posterior trajectories. We performed all analyses on trajectories pooled from all embryos imaged for a given condition and used bootstrap sampling of the pooled data to construct confidence intervals for each measurement. In all cases, analyzing data per embryo produced similar results.

### Measuring recruitment ratios and dissociation rates

To estimate recruitment ratios and rate constants for dissociation, we constructed single molecule release curves from the trajectories for anterior or posterior molecules as cumulative histograms *H*(τ_*n*_) defined as the number of trajectories with lifetime > τ_*n*_ = *n* × 50*ms*, *n* = 1,2, ….

To avoid counting false positives – e.g. cytoplasmic molecules that enter and leave the imaging plane without binding, we only considered trajectories with lifetimes >= 200 ms. We used Matlab’s non-linear least squares fitting function ***nlinfit*** to fit these release curves to a weighted sum of two exponential functions:

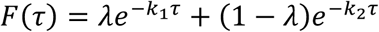

On the interval 200*ms* ≤ τ < ∞. We then corrected for photobleaching to estimate fast and slow dissociation rates:

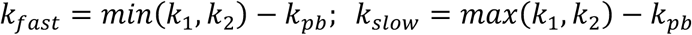

using the photobleaching rate *k_pb_* measured as above using the same imaging conditions and identical unit exposures. We computed the average dissociation rate for all molecules as the inverse of an average lifetime, corrected for photobleaching:

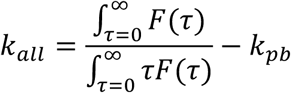

To estimate recruitment ratios from these fits, we took F(τ = 0) as our estimate of the total number of molecules binding to the anterior or posterior cortex (excluding false positives), and normalized by the total area of ROIs in which anterior or posterior molecules were observed to estimate the total number of molecules binding the anterior or posterior membrane per unit area and time:

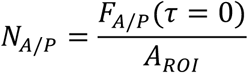

Then we took *NA/NP* as our estimate of the recruitment ratio.

### Measuring mobilities

To quantify single molecule mobilities, for all trajectories with lifetime >= 200 msec, we computed the distribution of displacements for a fixed interval 100 msec (D_100), measured for all segments of all trajectories pooled over all embryos. To compute reference distributions for PAR-3 oligomers, as shown in Figure 4C, we imaged endogenously-tagged PAR-3::GFP using the same acquisition parameters described above for single molecules, but with lower laser power (10%), and performed tracking analysis and constructed D_100 distributions as described above. To construct reference distributions for simulated Brownian diffusion, as shown in Figure 4C, we simulated 10,000 particles undergoing Brownian diffusion with D = 0.1 µm^2^/sec, added gaussian noise with mean 0 and SD = 50 nm to approximate localization errors in our system ^15^, then performed tracking analysis and constructed D_100 distributions as described above. We then used the linear least squares method to fit the D_100 distributions for single molecules to a weighted sum of the distributions for oligomers and simulated diffusion.

### PAR-3 depletion experiments

To obtain partial depletion of PAR-3, we placed young adult worms expressing endogenously-tagged PAR-3::GFP on RNAi feeding plates for 2-8 hours. To assess whether PAR-3 polarity was established prior to the onset of maintenance phase and whether posterior-directed cortical flows redistribute PAR-3 oligomers during maintenance phase, we imaged embryos in timelapse mode using 50 msec exposures and 30% laser power at 2.5-5 second intervals for 4 minutes following pseudocleavage relaxation. Immediately following this period, we imaged the same embryos in streaming mode with 100% laser power and 50 msec exposures for 50 seconds, and performed feature detection and oligomer size inference as described above. We determined the fluorescence density of segmented PAR-3 features in anterior and posterior ROIs and computed the anterior-posterior asymmetry of PAR-3 as the ratio of these values.

