## Supplementary material for "Oligomerization and positive feedback on membrane recruitment encode dynamically stable PAR-3 asymmetries in the *C. elegans* zygote": Modeling Supplement: biorxiv_version_Lang2025_suppl.pdf

In this section we describe the derivation, parameterization and analysis of a mathematical model for PAR-3 kinetics. The basic approach follows our previous theoretical analysis of oligomerization by peripheral membrane binding proteins [1]. Briefly, as schematized in Figs. 3A and 7A, we assume a well-mixed cytoplasmic pool of PAR-3 monomers which bind reversibly to the plasma membrane. At the membrane, monomers assemble reversibly into linear oligomers with size-dependent mobilities and dissociation rates. Finally, we assume that membrane-bound PAR-3 feeds back indirectly to promote the attachment of PAR-3 monomers to the membrane. Letting $A_n(x)$ be the density (number/length) of membrane-bound oligomers of size $n$ at point $x$, and $A_c$ be the concentration of cytoplasmic PAR-3, the density of membrane-bound oligomers evolves according to the following equations [1, Eq. (1)]

$$\partial_t A_1 = D_1 \partial_x^2 A_1 + \left(k_{\text{basal}} + k_f f(A)\right) A_c - k_{\text{off}} A_1 \qquad \text{(M1a)}$$
$$+ 2k_{\text{dp}} A_2 - 2k_{\text{p}} A_1^2 + \sum_{n=3}^{N} \left(k_{\text{dp}} A_n - k_{\text{p}} A_1 A_{n-1}\right)$$

$$\partial_t A_n = k_{\text{p}} A_1 \left(A_{n-1} - A_n\right) - k_{\text{dp}} \left(A_n - A_{n+1}\right) \qquad N > n \geq 2 \qquad \text{(M1b)}$$

$$\partial_t A_N = k_{\text{p}} A_1 A_{N-1} - k_{\text{dp}} A_N \qquad \text{(M1c)}$$

$$A_c = \frac{1}{hL}\left(A^{(\text{Tot})} L - \int_0^L A(x)\, dx\right) \qquad A(x) = \sum_{n=1}^{N} n A_n(x). \qquad \text{(M1d)}$$

The first line of (M1a) describes the membrane attachment and detachment of PAR-3 monomers, with basal rates governed by $k_{\text{basal}}$ and $k_{\text{off}}$, and density-dependent feedback on monomer attachment governed by $k_f f(A)$, whose form we consider below. The second line of (M1a), combined with (M1b) and (M1c), describes the dynamics of polymerization on the membrane, with forward and reverse rate constants $k_{\text{p}}$ and $k_{\text{dp}}$. Finally, the cytoplasmic concentration $A_c$ is determined in (M1d) from the conservation of total protein in the system, where $A(x)$ is the total bound density at each point on the membrane and $A^{(\text{Tot})}$ is the uniform density when all PAR-3 is bound.

The forms of equations (M1) reflect several important simplifying assumptions, which are justified by our experimental observations. First, our experimental data (Fig. 3K) show that oligomer mobilities and dissociation rates fall sharply with increasing oligomer size. Therefore we neglect both oligomer diffusion and detachment, as in [1]. Second, our experimental observations (see Section 1.1.2) suggest that direct attachment of cytoplasmic monomers to membrane-bound PAR-3 oligomers makes a relatively small contribution to the total flux of PAR-3 monomers onto the membrane. In previous work [1], we showed that direct attachment effectively reduces or increases the strength of indirect feedback, depending on the regime considered. Thus we do not explicitly model the direct attachment of cytoplasmic monomers to membrane-bound oligomers, instead assuming that the weak effects of this attachment are implicit in the feedback strength.

### 1 Nondimensionalization, analysis, and parameter estimation

To estimate the model parameters and compare model predictions to experimental observations, we scale the equations by appropriate density ($A^{(\text{Tot,WT})}$, which represents $A^{(\text{Tot})}$ under wild type conditions), length ($L$), and time ($1/k_{\text{dp}}$) scales. Since polymerization dynamics are much slower than detachment, ($1/k_{\text{dp}}$) is the time a typical PAR-3 molecule spends on the membrane. Accordingly, we define the dimensionless (hatted) variables

$$x = \hat{x}L, \qquad t = \hat{t}/k_{\text{dp}}, \qquad A = \hat{A}A^{(\text{Tot,WT})}.$$

Substituting into (M1) gives dynamics in dimensionless form [1, Eq. 14]

$$\begin{aligned}
\partial_{\hat{t}}\hat{A}_1 &= \hat{D}_1\partial_{\hat{x}}^2\hat{A}_1 + \hat{K}_{\text{on}}\left(1 + \hat{K}_{\text{f}}\hat{F}(\hat{A})\right)\hat{A}_{\text{c}} - \hat{K}_{\text{off}}\hat{A}_1 \\
&\quad + 2\hat{A}_2 - 2\hat{K}_{\text{p}}\hat{A}_1^{\,2} + \sum_{n=3}^{N}\left(\hat{A}_n - \hat{K}_{\text{p}}\hat{A}_1\hat{A}_{n-1}\right) && \text{(M2a)} \\
\partial_{\hat{t}}\hat{A}_n &= \hat{K}_{\text{p}}\hat{A}_1(\hat{A}_{n-1} - \hat{A}_n) - (\hat{A}_n - \hat{A}_{n+1}) \qquad N > n \geq 2 && \text{(M2b)} \\
\partial_t\hat{A}_N &= \hat{K}_{\text{p}}\hat{A}_1\hat{A}_{N-1} - \hat{A}_N && \text{(M2c)} \\
\hat{A}_{\text{c}} &= \hat{A}^{(\text{Tot})} - \int_0^1 \hat{A}(x)\,d\hat{x}, && \text{(M2d)}
\end{aligned}$$

Equations (M2) involve six dimensionless parameters

$$\hat{D}_1 = \frac{D_1}{L^2k_{\text{dp}}}, \quad \hat{K}_{\text{on}} = \frac{k_{\text{basal}}}{k_{\text{dp}}h}, \quad \hat{K}_{\text{f}} = \frac{k_fA^{(\text{Tot,WT})}}{k_{\text{basal}}}, \qquad \text{(M2e)}$$

$$\hat{K}_{\text{off}} = \frac{k_{\text{off}}}{k_{\text{dp}}}, \quad \hat{K}_{\text{p}} = \frac{k_{\text{p}}A^{(\text{Tot,WT})}}{k_{\text{dp}}}, \quad \hat{A}^{(\text{Tot})} = \frac{A^{(\text{Tot})}}{A^{(\text{Tot,WT})}}, \quad \hat{F}(\hat{A}) = \frac{f(A)}{A^{(\text{Tot,WT})}}.$$

(and the feedback function $F(\hat{A})$; see below), whose values we determine by direct measurement or by fitting to experimental data. Our data directly give values for three dimensional parameters:

the monomeric diffusion coefficient $D_1 = 0.1\ \mu\text{m}^2/\text{s}$, the monomer detachment rate $k_{\text{off}} = 3/\text{s}$, and the rate at which single subunits unbind from PAR-3 oligomers ($k_{\text{diss}} = 0.08/\text{s}$), which gives $k_{\text{dp}} = 0.16/\text{s}$ (since unbinding can occur from either side). We take the length scale $L = 134.6$ $\mu$m to be the perimeter length of the maximal cross-section of an ellipsoid, with half-axis lengths $27 \times 15 \times 15\ \mu$m, which approximates the shape and dimensions of a typical *C. elegans* embryo in mitosis [2]. Together, these measurements determine values for the non-dimensional parameters $\hat{D}_1$ and $\hat{K}_{\text{off}}$. The parameter $\hat{A}^{(\text{Tot})}$ represents a depletion factor; it is 1 in wild-type embryos and later varied to mimic depletion of PAR-3.

#### 1.1 Oligomerization and effective dissociation

To perform the analysis in Fig. 3(B–D), we first solve (M2) at steady state to obtain an exponential distribution of oligomer sizes [3, 1]

$$\hat{A}_n = \hat{K}_{\text{p}} \hat{A}_1 \hat{A}_{n-1} := \alpha \hat{A}_{n-1} \quad n \geq 2. \tag{M3}$$

This defines $\alpha = \hat{K}_{\text{p}} \hat{A}_1 = (k_{\text{p}}/k_{\text{dp}})\, A_1$ as the coefficient of the exponential distribution of oligomer sizes. It follows that the total amount of PAR-3 is given by

$$\hat{A} = \sum_{n=1}^{N} n \alpha^{n-1} \hat{A}_1 \xrightarrow{N \rightarrow \infty} \frac{\hat{A}_1}{(1-\alpha)^2}. \tag{M4}$$

This equation can then be solved for $\hat{A}_1$ to obtain [1, Eq. (12)]

$$\hat{A}_1 = \frac{1 + 2\hat{A}\hat{K}_{\text{p}} - \sqrt{1 + 4\hat{K}_{\text{p}}\hat{A}}}{2\hat{A}\left(\hat{K}_{\text{p}}\right)^2}, \tag{M5}$$

which gives the number of bound monomers as a function[1] of the total bound $\hat{A}$. The equations for $\alpha = \hat{K}_{\text{p}} \hat{A}_1$ and mean oligomer size $s = 1/(1-\alpha)$ given in the main text (and plotted in Fig. 3C) follow from (M5).

[1] This convenient closed-form expression is a consequence of our neglect of dimer and trimer unbinding. If dimer and trimer unbinding are included (as suggested by Fig. 3K), then (M3) only holds for $n \geq 4$, and $\hat{A}_1$ can be obtained from $\hat{A}$ by solving a quartic equation. The resulting model has negligible quantitative difference (at most 20% in the predicted recruitment asymmetry in polarized states) from the one we present here, which is significantly easier to analyze.

##### 1.1.1 Effective dissociation rate

In the case when $\alpha$ is known, we can compute the effective dissociation rate (Fig. 3D) by assuming that the distribution of oligomer sizes is given by (M3), with detachment rates given by $k_{\mathrm{off}}^{(n)} = k_{\mathrm{off}}^{(1)}\beta^{n-1}$. This gives the effective dissociation rate

$$k_{\mathrm{eff}} = \frac{k_{\mathrm{off}}^{(1)}\hat{A}_1}{\hat{A}}\sum_{n=1}^{\infty} n\alpha^{n-1}\beta^{n-1} = \frac{k_{\mathrm{off}}^{(1)}\hat{A}_1}{\hat{A}\,(1-\alpha\beta)^2} = k_{\mathrm{off}}^{(1)}\left(\frac{1-\alpha}{1-\alpha\beta}\right)^2, \tag{M6}$$

where $\alpha(\hat{A}) = \hat{A}_1(\hat{A})\hat{K}_{\mathrm{p}}$ is obtained using (M3) again. This results in the curve shown in Fig. 3D; note however that using an exponential relationship for the oligomer sizes and a nonzero detachment rate for oligomer sizes larger than one is inconsistent at steady state. The A/P ratio of 3.8 in the effective dissociation rate is obtained by substituting $\alpha = 0.73$ for the anterior and $\alpha = 0.42$ for the posterior into (M6), with $\beta = 1/4$ (fit from Fig. 3K).

##### 1.1.2 Estimating an upper bound on direct monomer binding fraction

To estimate an upper bound on the fraction of monomers that bind directly to membrane-bound oligomers, we assume energy conservation. Under this assumption, the fluxes of monomers on and off the membrane, and the fluxes of monomers in and out of oligomers, must both balance at steady state. The total flux of monomers onto the membrane is therefore

$$\delta_{\mathrm{mem}}(\hat{A}) := k_{\mathrm{off}}^{(1)}\hat{A}_1 \tag{M7}$$

while the total flux due to attachment of monomers to oligomers is given by

$$\delta_{\mathrm{olig}}(\hat{A}) := k_{\mathrm{dp}}\left(2\hat{A}_2 + \sum_{n=3}^{\infty}\hat{A}_n\right) = k_{\mathrm{dp}}\hat{A}_1\left(2\alpha + \sum_{n=3}^{\infty}\alpha^{n-1}\right) = \frac{\hat{A}_1 k_{\mathrm{dp}}(2-\alpha)\alpha}{1-\alpha} \tag{M8}$$

If all monomers that bind oligomers come from the cytoplasm, then the fraction of cytoplasmic monomers that bind directly to oligomers is given by

$$\frac{\delta_{\mathrm{olig}}(\hat{A})}{\delta_{\mathrm{mem}}(\hat{A})} = \frac{k_{\mathrm{dp}}(2-\alpha)\alpha}{k_{\mathrm{off}}^{(1)}\,(1-\alpha)} \tag{M9}$$

Using empirically measured values for $\alpha$ and the values for $k_{\mathrm{dp}} = 0.16/\mathrm{s}$ and $k_{\mathrm{off}}^{(1)} = 3/\mathrm{s}$ from Table M1, we estimate this fraction to be $\approx 0.18$ for anterior monomers ($\alpha = 0.73$) and $\approx 0.06$ for posterior monomers ($\alpha = 0.42$). However, because a fraction of the oligomer attachment flux must come from membrane-bound monomers, the real values are likely to be considerably lower.

##### 1.1.3 Simulations of detachment kinetics

To validate the model of polymerization kinetics against total intensity data, we simulate the equations (M2) without a spatial dimension, setting the polymerization rate $\hat{K}_\mathrm{p}$ according to our fit in Section 1.3. With $\hat{K}_\mathrm{p}$ fixed, we initialize $\alpha = 0.75$ on the anterior (slightly higher than in late maintenance phase), which sets the initial monomer concentration and total concentration. The monomer binding rate $\hat{K}_\mathrm{on}\left(1+\hat{K}_\mathrm{f}\hat{F}(\hat{A})\right)\hat{A}_\mathrm{c} = 0.201$ is held at a constant value, set so that the total PAR-3 at steady state is roughly 5/9 of the initial concentration (matching the end/start ratio in Fig. 4I). The resulting simulated curves are compared to the experimental curves in Fig. 4I (for total bound protein $\hat{A}$)

#### 1.2 Steady state analysis with feedback

Incorporating positive feedback due to recruitment, we first must choose the feedback function $f(A)$. Our experimental finding that PAR-3 recruitment asymmetries do not change during maintenance phase as levels of membrane-bound PAR-3 fall by $\approx 50\%$ suggest that positive feedback saturates at or below the density observed in late maintenance. Thus, we consider a simple model for saturating feedback given by

$$f(A) = \min\left(A, A^{(\mathrm{Sat})}\right) \to \hat{F}(\hat{A}) = \min\left(\hat{A}, \hat{A}^{(\mathrm{Sat})}\right), \tag{M10}$$

where $\hat{A}^{(\mathrm{Sat})} < \hat{A}_u$, the steady state bound PAR-3 density in the uniform state.

We first analyze the model behavior in the absence of diffusion, as experiments and simulations show that the boundary position in a polarized state is stable on timescales relevant to maintenance phase. Invoking the steady state (M3) for polymerization dynamics reduces the monomer equation (M2a) to the balance of an attachment flux $\delta(\hat{A})$ (which sums the fluxes due to basal attachment and feedback) and the detachment flux $\nu(\hat{A})$,

$$0 = \underbrace{\hat{K}_\mathrm{on}\left(1+\hat{K}_\mathrm{f}\hat{F}^{+}(\hat{A})\right)\hat{A}_\mathrm{c}}_{\text{Attachment, }\delta(\hat{A})} - \underbrace{\hat{K}_\mathrm{off}\hat{A}_1(\hat{A})}_{\text{Detachment, }\nu(\hat{A})} \tag{M11}$$

where $\hat{A}_1(\hat{A})$ is given in (M5). At spatially uniform steady states, $\hat{A}(x) \equiv \hat{A}$, the dimensionless cytoplasmic concentration (defined in (M2d)) becomes $\hat{A}_\mathrm{c} = \hat{A}^{(\mathrm{Tot})} - \hat{A}$ (recall $\hat{A}^{(\mathrm{Tot})} = 1$ for wild type embryos), and the total attachment rate is

$$\delta(\hat{A}) = \delta_\mathrm{cons}(\hat{A}) = \hat{K}_\mathrm{on}\left(1+\hat{K}_\mathrm{f}\hat{F}^{+}(\hat{A})\right)\left(1-\hat{A}\right) \tag{M12}$$

In this case, the attachment and detachment curves in (M11) depend only on $\hat{A}$, and spatially uniform steady state(s) can be read from their intersections on a flux balance plot (solid red and dotted blue lines in Fig. M1). Following the Local Perturbation Analysis (LPA) approach of [4], we determine stability by considering the response to a local perturbation $\hat{A} = \hat{A}_u + \epsilon\hat{A}$ of finite size within an infinitesimally small domain such that the cytoplasmic concentration remains constant ($\hat{A}_\text{c} = 1 - \hat{A}_u$). This situation corresponds to an attachment rate of

$$\delta(\hat{A}) = \delta_\text{LPA}(\hat{A}) = \hat{K}_\text{on}\left(1 + \hat{K}_\text{f}\hat{F}^+(\hat{A})\right)\left(1 - \hat{A}_u\right). \tag{M13}$$

The qualitative response of the system can be inferred by examining the behavior of $\Delta(\hat{A}) = \delta_\text{LPA}(\hat{A}) - \nu(\hat{A})$. The case when $\Delta'(\hat{A}_u) > 0$ corresponds to an unstable uniform state, while $\Delta'(\hat{A}_u) < 0$ corresponds to a stable one.

We use the saturated feedback function (M10) to identify spatially uniform steady states as solutions of the flux balance equation (M11) with $\delta = \delta_\text{cons}(\hat{A})$ given in (M12). We then evaluate the possibility for inducible polarity using LPA; as shown in the flux balance plot in Fig. M1, if the feedback saturates *below* the spatially uniform steady state concentration, the spatially uniform steady state is always locally stable, since $\delta'_\text{LPA} = 0$ if saturated with $\nu' > 0$. For a subset of values of ($\hat{K}_\text{on}$, $\hat{K}_\text{f}$ and $\hat{A}^{(\text{Sat})}$), a transition to a stably polarized state can be induced by local perturbations that drive $\hat{A}$ *below* some threshold, as indicated by the second crossing of the fluxes in Fig. M1. In this case, at the stably polarized steady state, $\hat{A}_\ell$ lies below the threshold for inducing polarity, while $\hat{A}_h$ is held close to spatially uniform density $\hat{A}_u$ by saturating feedback. As discussed in the main text, this is the scenario which agrees best with experimental observations.

This analysis gives qualitative constraints on the feedback saturation level $\hat{A}^{(\text{Sat})}$. If $\hat{A}^{(\text{Sat})} > \hat{A}_u$, the uniform state could be unstable, as $\delta'_\text{LPA} > 0$ if the feedback is not saturated. On the other hand, if $\hat{A}^{(\text{Sat})}$ is too small, an asymmetry induced by feedback will be impossible. We will make the (arbitrary) assumption that $\hat{A}^{(\text{Sat})} = 0.8\hat{A}_u$, noting that our results are not sensitive to the precise value used.

##### 1.3 Phase diagram and parameter estimation

Having analyzed the model, we are now in position to assign parameters based on experimental observations. To fit the remaining dimensionless parameters[2] $\hat{K}_\text{on}$, $\hat{K}_\text{f}$, and $\hat{K}_\text{p}$, we use the following

[2] The parameter $h$ (cytoplasmic "thickness") only appears in the dimensionless parameter $\hat{K}_\text{on}$. Since we fit $\hat{K}_\text{on}$, we never assign a value to $h$ in Table M1.

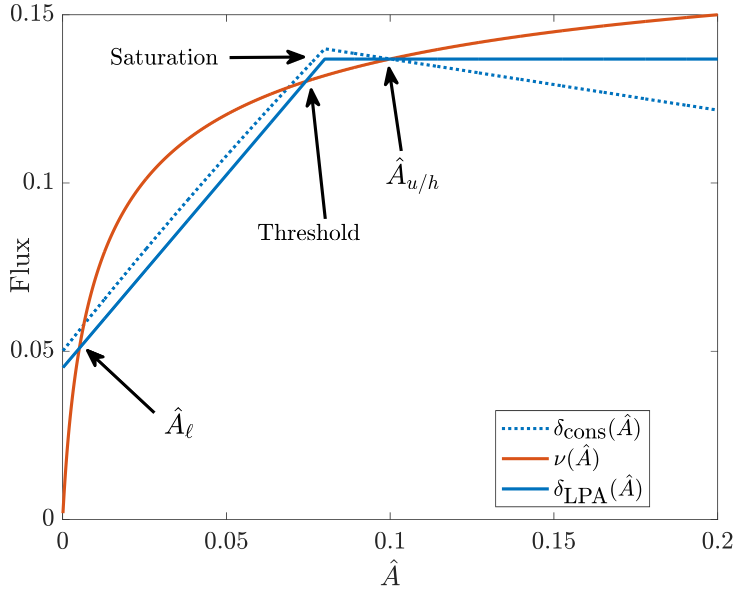


Figure M1: Flux plane analysis for saturated feedback model (M10). Given $\hat{A}^{(\text{Sat})}$, the uniform state is defined by the crossing of the attachment rate $\delta_{\text{cons}}(\hat{A})$ (dotted blue lines, see (M12)) with the detachment rate $\nu(\hat{A})$ (solid red lines). *Stability* of the uniform state is determined by how the attachment rate $\delta_{\text{LPA}}(\hat{A})$ (solid blue lines, see (M13)) compares to $\nu(\hat{A})$ near the steady state. The uniform state is stable if the feedback saturates below it.

| Parameter | Description | Value | Units | Ref | Notes |
|---|---|---|---|---|---|
| $L$ | Domain length | 134.6 | $\mu$m | [2] | radii $27 \times 15$ $\mu$m ellipse |
| $D_1$ | Monomeric PAR-3 diffusivity | 0.1 | $\mu\text{m}^2/\text{s}$ | | This study |
| $\hat{K}_{\text{on}}$ | Monomeric PAR-3 attachment rate | 0.10 | | | Assume 10% bound PAR-3 |
| $\hat{K}_{\text{f}}$ | Feedback strength | 14.5 | | | Fit recruitment asymmetry |
| $\hat{A}_u$ | Uniform state PAR-3 level | 0.10 | | | Photobleaching experiments |
| $\hat{A}^{(\text{Sat})}$ | Feedback saturation level | 0.08 | | | 80% of uniform state in wild type |
| $k_{\text{off}}$ | Monomeric PAR-3 detachment rate | 3 | 1/s | | This study, Fig. 3K |
| $k_{\text{dp}}$ | PAR-3 depolymerization rate | 0.16 | 1/s | | This study, Fig. 4 |
| $\hat{K}_{\text{p}}$ | PAR-3 polymerization rate | 62 | | | Fit mean oligomer size |
| $N$ | Max oligomer size | 50 | | | Same results for larger $N$ |

Table M1: Parameter values for the PAR-3 model (see (M2e) for the definitions of dimensionless parameters).

procedure:

1. Based on photobleaching experiments, we estimate that roughly 10% of the total pool of PAR-3 is bound to the membrane at steady state. Thus, for every choice of $\hat{K}_{\text{f}}$ and $\hat{K}_{\text{p}}$, we fix $\hat{K}_{\text{on}}$ so that 10% of protein is bound to the cortex at the uniform steady state.

2. We scan over pairs of parameters $(\hat{K}_{\text{p}}, \hat{K}_{\text{f}})$ and use LPA to produce a phase diagram of where a polarized state coexists with the spatially uniform state (Fig. 6B).

3. For pairs of points where polarization is theoretically possible, we run simulations of the polarized state. Our initial conditions correspond to the end of establishment phase: we use a 10:1 A/P asymmetry, with 50% of the domain considered anterior, and assume local equilibrium of oligomerization reactions with 15% of the total protein pool bound to the membrane (this is reduced by $\sim 40\%$ during maintenance phase). Evolving this initial state for four minutes, we obtain an anterior mean oligomer size

$$\text{Mean oligomer size} = \left(\sum_{n=1}^{N} n\hat{A}_n\right) / \left(\sum_{n=1}^{N} \hat{A}_n\right). \tag{M14}$$

   and recruitment asymmetry $\max(\delta(\hat{x}))/\min(\delta(\hat{x}))$, which we plot in Fig. 6C.

4. The correct values of $\hat{K}_{\text{p}} = 62$ and $\hat{K}_{\text{f}} = 14.5$ are obtained by matching the anterior mean oligomer size and recruitment asymmetry to experimental data. The model predicts a correspond A/P asymmetry of $\hat{A}_h/\hat{A}_\ell = 8$, also in agreement with the data.

This completes parameter selection, which is summarized in Table M1.

### 2 Dynamics

We use standard numerical methods to solve (M2), discretizing the one-dimensional domain at $N$ points, with diffusion being discretized via the standard three point Laplacian. We use a forward Euler method for the reaction terms, and a backward Euler method for the diffusion terms (in the monomer equation), which gives first order accuracy in time [5, c. 1]. Typical parameters are $N = 1000$ (required to resolve the boundary layer where the enriched PAR-3 domain can shift) and $\Delta\hat{t} = 0.02$ (50 steps per depolymerization time). Matlab code is available at the github repository `https://github.com/omaxian/CElegansModel/`.

We first simulate our spatial PAR-3 model with two different initial conditions (Fig. M2): a perturbation of the uniform state (which demonstrates that the uniform state is a stable attractor)

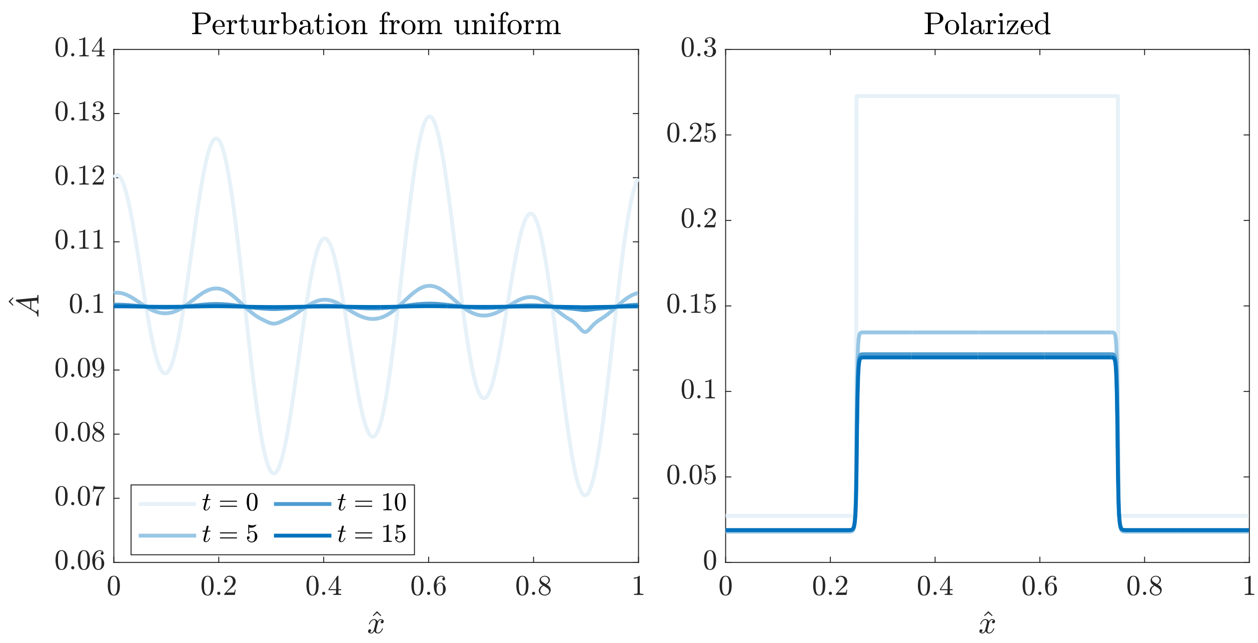


Figure M2: Dynamics of the saturated feedback model. At left, we show that perturbations to the uniform state decay, while at right, we show that the polarized state is stable in time. Dynamics occur on timescales of 5–10 minutes (legend).

and the polarized state starting at the end of establishment phase, which decays to a stable polarized state. In both cases, the timescale on which the protein levels adjust is roughly 5–10 minutes. We notice barely any drift of the boundary on this timescale, which supports our neglect of diffusion in model analysis.

##### 2.1 Systematic depletion of PAR-3

To model the experiment of PAR-3 depletion (Fig. 6D), we vary the total fraction $\hat{A}^{(\text{Tot})}$ of PAR-3 in (M2d). To simulate experimental conditions for a given $\hat{A}^{(\text{Tot})}$, we start at the end of establishment (see Section 1.3), then run until we reach $t = 4$ minutes, which corresponds to the time interval used in experiments. Because the dimensionless parameters in (M2e) are in terms of the total PAR-3 in *wild-type* embryos, only $\hat{A}^{(\text{Tot})}$ changes when we perform the depletion experiment.

The results in Fig. 3I show that our model successfully reproduces the trend in mean oligomer size vs. total bound density (measured on the anterior at $t = 4$ mins). For the A/P asymmetries in Fig. 6D, our model captures the qualitative and quantitative trend of no asymmetry for low mean oligomer sizes, followed by a transition to stable asymmetries on the order 6–8 for larger mean oligomer sizes. This is in contrast to a model based on asymmetric recruitment (in which case there is no feedback, the basal rate is 26% higher, and monomers are recruited to the anterior

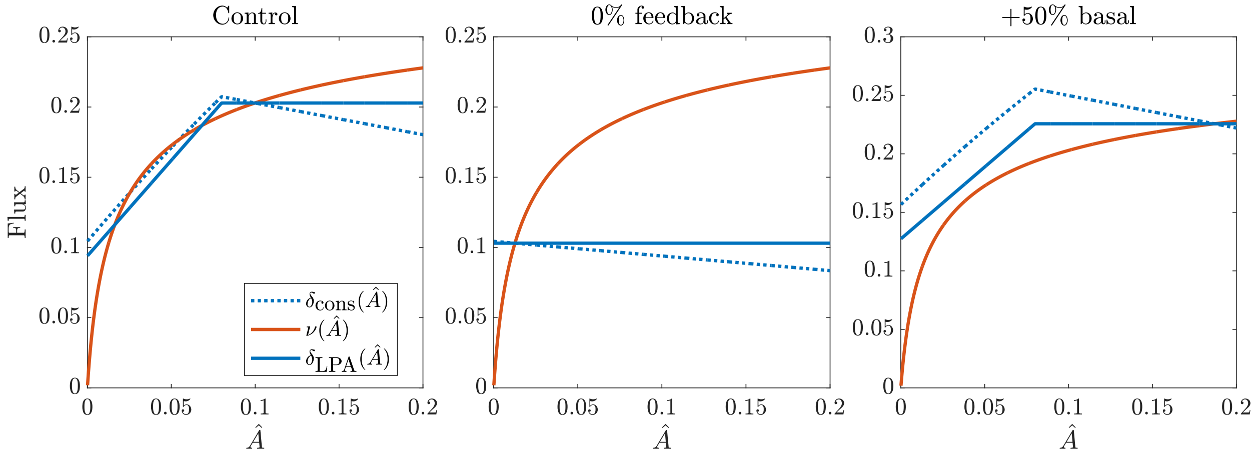


Figure M3: How the system responds to reductions in feedback and changes in the basal rate. Here we show theoretical flux balance plots when considering default parameters ($\hat{K}_{\text{f}} = 14.5$, $\hat{K}_{\text{on}} = 0.10$), 0% feedback ($\hat{K}_{\text{f}} = 0$, $\hat{K}_{\text{on}} = 0.10$), and 150% basal binding rate ($\hat{K}_{\text{f}} = 9.7$, $\hat{K}_{\text{on}} = 0.15$). The main text (Fig. 8G) shows the corresponding simulated polarized states.

at 1.72 times the rate of the posterior), which predicts a persistent asymmetry even as the mean oligomer size drops to 1 (i.e., even without oligomerization). Note that the two curves overlap (for the second time) at the "wild type" parameters.

#### 2.2 Changing feedback to mimic aPAR depletion

Finally, we use our saturated feedback model to demonstrate what might happen when the feedback is reduced, in order to explain the results of some of our perturbation experiments. To do this, we perturb the feedback parameter $\hat{K}_{\text{f}}$ and examine changes to the flux balance plot (Fig. M3) and dynamics of an initially polarized state (10:1 asymmetry with 50% protein bound; main text Fig. 8). We are already familiar with the flux balance plot for stably polarized states (Fig. M1). Simulating maintenance phase with control parameters yields the expected dynamics: the anterior domain concentration decreases, while the posterior remains roughly constant, and there is a steady state asymmetry.

Reducing to 0% feedback ($\hat{K}_{\text{f}} = 0$) gives dynamics which correspond roughly to our experiment with PAR-6 depletion (RNAi). In this case, removing feedback gives a single, smaller, uniform steady state which is close to posterior levels in the wild type (middle panel of Fig. M3). In simulations (Fig. 8G), we consequently observe a convergence of the anterior and posterior levels onto a smaller concentration, near to the original posterior level.

For CDC-42 depletion (RNAi), we hypothesize that an increase in the basal binding rate could lead to a loss of asymmetry. To test this, we increase $k_{\text{basal}}$ by 50%; using the non-dimensionalization in (M2), this implies $\hat{K}_{\text{on}}$ increases by 50% and $\hat{K}_{\text{f}}$ decreases by 33%. As shown in the right panel in Fig. M3, the effect of this change is to increase the $y$ intercept of the local flux balance curve $\delta_{\text{LPA}}$, which leads to a loss of bistability. The dynamics with this choice of parameters roughly correspond to our experiment in CDC-42 (RNAi) embryos, where the posterior state rises up to meet the anterior one (Fig. 8G).
